# Single-cell transcriptome of retinal myeloid cells in response to transplantation of human neurons reveals reversibility of microglial activation

**DOI:** 10.1101/2025.05.16.654622

**Authors:** Emil Kriukov, Anthony Mukwaya, Paul Francis Cullen, George Baldwin, Volha V. Malechka, Nasrin Refaian, Nikita Bagaev, Everett Labrecque, Sthavir Vinjamuri, Milica A. Margeta, Petr Baranov

**Affiliations:** Schepens Eye Research Institute of Massachusetts Eye and Ear, Boston, MA, USA; Department of Ophthalmology, Harvard Medical School, Boston, MA, USA

## Abstract

The host retinal microglia and macrophage activation remains a major challenge for the integration of donor neurons following transplantation. Previously, we and others have shown that it is possible to increase donor retinal ganglion cell (RGC) survival by inhibiting the microglia-RGC interaction with Annexin V or through reprogramming microglia with the soluble Fas ligand. However, the exact mechanisms of the microglia/macrophage activation and their heterogeneity following transplantation remain unknown. To address this question, the donor RGC were differentiated from Brn3b-Tdtomato-Thy1.2 human embryonic stem cells using a 3D protocol, followed by dissociation and RGC purification. RGC were delivered subretinally (1.5x10^4^ viable cells/eye) into 3-6-month-old *CX3CR1^GFP^*knock-in mice. Three days after transplantation retinas were dissociated into single-cell suspension and GFP-positive myeloid cells isolated using FACS. Of the sorted cells, up to 10,000 viable cells per sample were used for single-cell RNA library preparation and sequenced using the 10X Genomics Chromium platform. In addition, several retinas were fixed and stained for donor RGC (mCherry) and host microglia/macrophages (Iba1). RNA Velocity was used to reconstruct the myeloid cell population and activation trajectory from scRNAseq data. We observed continuous bi-directional transition of microglia/macrophages from a homeostatic to an activated state. We also observed that the response to the transplant falls into the classic disease-associated-microglia (DAM) activation paradigm with a decrease in expression of the homeostatic gene *Tmem119* and an increase in expression of disease-associated genes including *Apoe, Lgals3,* and *Spp1*. Our findings show that the host retinal myeloid cell population undergoes activation upon transplantation of stem-cell derived donor RGC, with a molecular profile of the activated cells similar to that of activated myeloid cells associated with neurodegenerative diseases of the brain and the eye. Advanced integrated transcriptomic analysis shows separate activated-to-homeostatic and homeostatic-to-activated trajectories suggesting the reversibility of this process.

## INTRODUCTION

Retinal ganglion cells (RGC) play a critical role in the transmission of visual signals from the retina to the brain. The optic neuropathies, including glaucoma, non-arteritic anterior ischemic optic neuropathy, Leber Hereditary Optic Neuropathy, traumatic optic neuropathy, and optic pathway glioma result in damage to RGC, ultimately leading to their death. RGC loss is permanent given that these cells cannot regenerate in mammals.^1^ Neuroregenerative strategies for targeting vision loss in optic neuropathies have been mainly focused on restoring the lost RGC cell population in a glaucomatous retina. This approach not only aims to halt disease progression, but also to restore the lost vision by replacing the damaged/lost RGC through RGC transplantation.^2–4^ To achieve functional regeneration, the transplanted RGC must survive, integrate, and extend their axons and establish connections within the retina and the brain. Of the many factors influencing initial donor RGC survival, the innate immunity^5^ and primarily microglia reactivity^6,7^ are of particular interest. In animal models of glaucoma, microglia have been heavily implicated in driving disease progression, promoting a pro-inflammatory environment. ^8,9^ In some instances, activated microglia have been directly implicated in RGC loss by mediating phagocytosis of the damaged or injured RGC.^10^ In line with this, our recent work shows that activated host microglia are detrimental to donor RGC survival, presumably via phagocytosis of the donor RGC. We observed that pre-treatment of donor RGC with sFasL and Annexin V to block the exposed phosphatidylserine residues of the stressed donor RGC prior to transplantation reduced activation of host microglia, and resulted in better RGC survival rates post transplantation.^11^ Studies like these and others underscore the importance of modulating host microglia reactivity for better transplantation outcomes.

To further improve donor RGC survival after transplantation, a better understanding of the host myeloid cell reactivity following donor RGC transplantation is paramount. We and others have previously shown the importance of modulating the retinal microenvironment to promote RGC survival.^2,4,12–16^ In the current study, we profiled the transcriptome of the host myeloid (*CX3CR1^GFP^*) cell population using a single-cell RNA sequencing approach, to delineate the myeloid cell responses on a single-cell level. A focused analysis of the microglial cell population was performed, given their established role in glaucoma disease progression, and compared to our healthy adult mouse retina cell atlas. We demonstrate that microglia following RGC transplantation upregulate disease associated microglia (DAM) genes, including *Apoe*, *Spp1* and *Lgals3*, which are classically seen in neurodegenerative diseases of the brain and the retina.^17–19^ In addition, we identified several other genes associated with microglia activation, including *Cybb*, *Atf3*, *Aurka* and *Kif22*. Given the known function of these genes in other disease contexts, we believe that these genes are important for modulating donor RGC survival post transplantation and thus, warrant further investigation in this context. Taken together, we present a comprehensive analysis of the host myeloid cell population following transplantation at a single-cell resolution level, specifically focusing on microglia reactivity, and provide insights into the regulatory mechanisms that may be important for donor RGC survival and integration into injured retina.

## MATERIALS AND METHODS

### Animals

*CX3CR1^GFP^* 3–6-month-old knock-in mice of mixed gender were purchased from Jackson Laboratory (Bar Harbor, ME). The animals were bred and maintained as *CX3CR1^GFP^* heterozygotes on C57BL6 genetic background at the Schepens Eye Research Institute (SERI). The animals were group-housed in a temperature-controlled environment with a standard 12-hour dark-light cycle with food and water provided *ad libitum*. All experimental procedures conformed to the National Research Council’s Guide for the Care and Use of Laboratory Animals, the Association for Research in Vision and Ophthalmology (ARVO) guidelines and were approved by the Institutional Animal Care and Use Committee (IACUC) of the SERI.

### Human stem cell-derived RGC differentiation

H9-BRN3B:tdTomatoThy1.2-human embryonic stem cells (hESCs) were maintained as colonies on six-well plates pre-coated with 2% Matrigel and grown in mTeSR1 medium under low oxygen conditions (3.5% O2, 5% CO2, 95% humidity, 37°C). Medium was changed daily and upon reaching approximately 70% confluency, hESCs were passaged using accutase for 8 to 12 minutes at room temperature and either split in the ratio of 1:10 to 1:20 or used for differentiation.

Human ESCs were differentiated into 3D retinal organoids as previously described. Briefly, colonies of hESCs were enzymatically dissociated for 12 minutes at room temperature into single cells using accutase. To initiate the formation of embryoid bodies (EBs), 7,500 hESCs were seeded per well of an ultra-low attachment U-bottom 96-well plate and cultured under low oxygen condition in mTeSR1 medium supplemented with 20 μM Y-27632. From day 1, differentiation medium was added daily and gradually transitioned from a 1:1 mix of mTeSR1 and BE6.2-NIM [B27 + E6 at 2x concentration – neural induction medium: 1x L-glutamine, 1x sodium pyruvate, 1x antibiotic-antimycotic, 1x B27 without Vitamin A, 0.88 mg/mL NaCl, 38.8 mg/L insulin, 128 mg/L L-ascorbic acid, 28 μg/L selenium, 21.4 mg/L transferrin, and 38.8 mg/L NaHCO3 in DMEM/F12 (3500 mg/L dextrose)] to primarily BE6.2-NIM by day 8. The differentiation medium was supplemented with 9 μM IWR-1-endo and 1% Matrigel from day 1 to day 4 and with 55 ng/mL BMP4 on day 6, to promote neural development early retinal differentiation. On day 8, organoids were transferred in a standard petri dish and randomly chopped to dissect the optic vesicles and to allow better propagation and greater expansion. To support attachment of organoids and growth of retinal epithelium, the chopped retinal aggregates were transferred in a tissue culture treated T75 flask in Long-Term Retinal Differentiation Medium (LT-RDM) [1x L-glutamine, 1x sodium pyruvate, 1x antibiotic-antimycotic, 1x B27, and 1x non-essential amino acids, 50:50 1x DMEM/F12 (3500 mg/L Dextrose): DMEM (4500 mg/mL dextrose)] supplemented with 10% FBS and 300 nM smoothened agonist (SAG), and cultured under normoxic conditions (20% O2, 5% CO2, 95% humidity, 37°C). LT-RDM supplemented with 300 nM SAG was replaced every other day with a gradually decreasing from 10% FBS to 0% FBS by day 15. On day 15, the clusters of retinal tissue were moved to an ultra-low attachment T75 flask for suspension culture in RDM supplemented with 300 nM SAG. The next day, 2/3 of medium was replaced with LT-RDM supplemented with 300 nM SAG and 2.5% FBS. On day 18, 2/3 of medium was replaced with LT-RDM supplemented with 300 nM SAG and 5% FBS. The medium was replaced every other day with LT-RDM supplemented with 7.5% FBS on day 20 and 10% FBS on day 22 to day 50. LT-RDM was supplemented with 1mM taurine from day 24 to 50, with 125 nM all-trans-retinoic acid (ATRA) on day 26, and with 250 nM ATRA from day 28 to day 50. Staring on day 34, LT-RDM was supplemented with 10 µM (2S)-N-[(3,5-Difluorophenyl)acetyl]-L-alanyl-2-phenyl]glycine 1,1-dimethylethyl ester (DAPT). The LT-RDM was replaced every other day until the retinal organoids were dissociated using the Miltenyi Biotec embryoid body dissociation kit according to the manufacturer protocol with the gentleMACS™ Dissociator protocol (Brain_01 and Brain_02).

Human RGC were sorted by magnetic microbead using the CD90.2 Dynabeads (Miltenyi Biotec, CD90.2 MicroBeads, mouse) according to the manufacturer protocol.

### *In vivo* transplantation of human ES-derived RGC

Mice were anesthetized intraperitoneally with 100 mg/kg ketamine hydrochloride and 20 mg/kg xylazine hydrochloride solution diluted with saline and allowed to be fully sedated. Pupils were dilated with 1% tropicamide ophthalmic solution (Baush & Lomb Americas Inc., Bridgewater, NJ). One drop of 0.5% proparacaine hydrochloride ophthalmic solution (Bausch & Lomb Americas Inc., Bridgewater, NJ) was applied onto the surface of the cornea for 2 minutes. Using a glass pipette, 1 μl solution containing 15,000 viable tdTomato positive human ES-derived RGC was injected into eye, one eye per mouse. One drop of Neomycin and Polymyxin B Sulfate and Bacitracin Zinc Ophthalmic ointment (Bausch & Lomb Americas Inc., Bridgewater, NJ) was applied to the surface of the injected eye following the procedure.

### Mouse retina isolation, dissociation, and single-cell suspension preparation

Three days after RGC transplantation, mice were euthanized, injected eyes enucleated and retinal tissue isolated and immediately transferred into a tube containing ice cold phosphate-buffered saline (PBS) with 0.04% bovine serum albumin (BSA). Eight mouse retinas were pooled to make a sample. The tissue was initially dissociated mechanically, followed by enzymatic method using the Gentle MACS protocol (Miltenyi Biotech, Adult Brain Dissociation Kit, mouse, and rat). Briefly, retinas were allowed to settle to the bottom of the tube and the excess PBS carefully removed with the help of a pipette. The tissues were then rinsed with Hanks’ balanced salt solution (HBSS) and transferred to a MACS M-tube and again left let to settle to the bottom of the tube. Excess HBSS was removed from the MACS tube with the aid of a pipette. The tissues were then resuspended in 2 mL of dissociation solution containing DNAse 1 (1 mg/mL). The tissue-enzyme mixture was incubated at 37°C for 5 minutes in a heating bath. The M-tube was then rapidly inverted ensuring that all dissociated tissues remained in solution. The tube was then placed into the MACS tissue dissociator and the brain 1 program initiated. At the end of the brain 1 program, the M-tube was removed from the MACS dissociator and incubated at 37°C for 5 minutes in a heating bath. The tube was then rapidly inverted and placed back into the MACS dissociator and program brain 2 initiated. At the end of the run, 10 mL of isolation buffer was added to the dissociate and the mixture triturated 10 times using a 1000 µL pipette. Then 12 mL of the homogenate was transferred into a 15 mL tube and cells pelleted at 200g for 5 minutes at 4°C. The resultant cell pellet was resuspended in 500 µl of isolation buffer and filtered through a 70u cell strainer into a FACS collection tube in preparation for cell for sorting.

### Fluorescent activated cell sorting (FACS)

All tubes were initially pre-blocked with 2% BSA in HBSS. Single stain samples for each labeled cell class, i.e. TdTomato and GFP+ were used to set the gates. Cells were sorted for TdTomato positive only, and GFP positive only, and collected into separate collection tubes which contained 200μl of PBS with 0.04% BSA. The sorted single-cell suspension was immediately used for library preparation for single-cell RNA sequencing.

### Single-cell RNA library preparation and sequencing

Up to 10,000 viable cells per sample were used for library preparation. Libraries were prepared following the 10x Genomics Chromium Next GEM Single Cell 3 Reagent Kits v3.1 protocol (Chromium Single Cell V(D) J Reagent Kits with Feature Barcoding technology for Cell Surface Protein, Document Number CG000186 Rev A, 10x Genomics, (2019, July 25). Briefly, the droplet-based encapsulation method was used to encapsulate the single cells with gel beads, partitioning oil, and a reverse transcription master mix generating gene expression emulsions (GEMs). The generated GEMs were cleaned, expression libraries constructed, barcoded and sequenced at a sequencing depth of 20,000 read pairs per cell, with paired-end dual indexing (See manufactures instructions for details).

### Quality control and data processing

Raw data was processed using Cell ranger v. 7.0.1. A mouse reference genome was used for sequence alignment (2024-A Mouse GRCm39). Cell ranger output that included matrix, features, and barcodes, was processed using Seurat v. 4.3.0.1.^20^ Quality control metrics included filtering by nCount_RNA, nFeature_RNA, percent.mt, percent.rb. Doublets were removed using the DoubletFinder algorithm.^21^

### Trajectory reconstruction with RNA Velocity

The output of the Cell ranger pipeline including possorted_genome_bam.bam files were used to generate .loom files using velocyto.^22^ Upon the concatenation of the .h5ad FA2-embedded dataset to the .loom splicing matrix, RNA Velocity analysis was performed using scVelo.^23^ The dynamical mode was applied to generate the velocity directions, then upon Leiden clustering the data was subdivided into a ‘homeostatic’ and ‘activated’ cell states based on the directionality of their velocity vectors. Downstream analysis to gain insight into regulatory mechanisms was performed using the two defined population states, i.e., homeostatic (Tmem119^+^) and activated (Apoe^+^) subsets.

### RNA Velocity-based commitment score estimation

To assess commitment of cells within the two identified cell states i.e. homeostatic and activated, and homeostatic populations towards the homeostatic and activated states, cell-level RNA velocity vectors were computed, with data obtained from an integrated single-cell RNA sequencing dataset. Generally, a cell’s velocity vector consists of two components x and y, representing direction and magnitude respectively in a 2D reduced-dimensional ForceAtlas2 embedding.

For each velocity vector *i*, the magnitude is computed as the Euclidean norm:

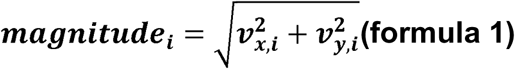

The direction of each vector *i* is described by an angle computed using the atan2 function:

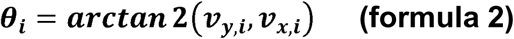

The angle is then converted from radians to degrees and standardized to the range [0, 360):

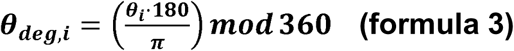

To assess directional bias, the range of angles (0–360 degrees) is discretized into bins. 36 bins of 10 degrees each were used. Once binned, the magnitudes of all velocity vectors falling into the same angular bin are summed. This yields a “cumulative magnitude” per direction bin.

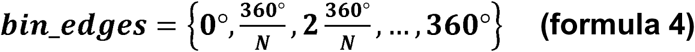

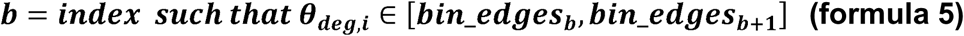

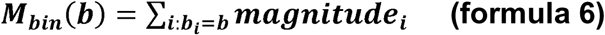

Two datasets (mg_tr1 for activated and mg_tr2 for homeostatic subsets) are processed similarly. Their directional cumulative magnitudes were computed. Ratios of cumulative magnitudes were calculated to quantify directional biases or differences in overall velocity distribution between conditions for the angular regions A and B based on the vector directionality. The resultant ratio represents unnormalized commitment score (unCS).

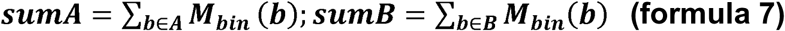

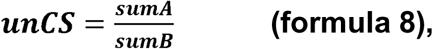

where unCS > 1 shows commitment of mg_tr1 towards homeostatic state

Since cell commitment depends on cumulative magnitude, which is highly dependent on the number of cells, further normalization is required. Cell numbers can be influenced by either the experimental condition or batch effect between samples or both. Given this, the real commitment score lays between the unnormalized and normalized commitment scores. Normalized commitment score (nCS) is quantified with the number of cells adjustment for populations 1 and 2 as:

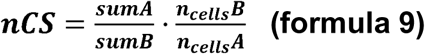

### Assembly of selected publicly available single-cell RNA sequencing datasets for the generation of a mouse reference atlas

To generate a representative mouse reference atlas for efficient mapping of myeloid cell populations from a range of different experimental conditions, we selected publicly available mouse datasets representing health and disease conditions of mice. For the health conditions, we focused on datasets generated from ‘developing mouse retina’, and ‘healthy adult mouse retina’. For the disease conditions, we focused on datasets generated from two mouse models of disease: ‘optic nerve crush model (ONC)’, and the ‘mouse microbead model of glaucoma’. A complete list of the selected datasets is summarized in **Supplementary Table 1**. Quality control and data processing was performed independently for each dataset, followed by data integration either by: 1) default Seurat integration, 2) Seurat RPCA-based integration, 3) Harmony^24^ or scVI^25^ . The method of choice for data integration was determined by the number of batches, batch variation, and by the total number of cells. To allow further analysis of myeloid between the conditions, we subset the healthy wild type myeloid population from every condition, as well as from the host upon transplantation myeloid dataset. Each was rescaled, and the subsequent integration was performed using default Seurat integration algorithm. To allow cell-cell interactions analysis and reference mapping of the host upon transplantation myeloid based on the mouse healthy adult retina atlas, we identified the cell classes present in the atlas using the differentially expressed genes and commonly known cell class markers.

### Healthy adult myeloid trajectory reconstruction

We sampled the myeloid population from the mouse healthy adult retina atlas. Given the number of batches involved, and variation in the number of cells per batch, we applied scVI to generate the mouse healthy adult myeloid subset embedding using 2 hidden layers and 30 latent spaces. Using the scVI-generated embedding, we performed cell fate trajectory reconstruction using scFates^26^ approach. With the reconstructed trajectory, we identified three states of the myeloid cell population. These were: 1) microglia/macrophage, 2) T cells, and 3) dendritic cells. This approach allowed for a more precise reference mapping of our experimental host upon transplantation myeloid population dataset, on to the defined microglia/macrophage myeloid population only. Annotation labels for myeloid cell population upon trajectory reconstruction were added to the healthy adult retina atlas, and the atlas was then used as a reference tool in Azimuth-based ^27^ annotation of our host upon transplantation myeloid cell population dataset.

### *In silico* myeloid replacement and cell-cell interactions analysis

To perform *in silico* myeloid replacement experiments, we used the mouse healthy adult retina to mimic the ‘recipient/host’. To perform the experiment, the compiled dataset was split into three developmental timepoints i.e. young adult, mature adult, and aged, based on the number of weeks postnatal. The software CellChat v. 2.1.0 ^28,29^ was used to analyze cell-cell communication, between the cells. The pipeline includes a few non-default settings: trim = 0.1, raw.use = F, population. Size = F, type = ‘truncatedMean” in computeCommunProb(), smoothData using PPI.mouse. Timepoints with low myeloid abundance were skipped to avoid false positives. The myeloid population of the mouse healthy adult retina was replaced in silico with either 1) host myeloid population upon transplantation, or 2) healthy developing retina myeloid cell population. In both cases, CellChat was used to analyze cell-cell interaction between cells. The resultant three sets of CellChat objects were then explored in a timepoint dependent manner. Using the 20 weeks old time point as proxy for a (6-month-old mouse, one into which RGC would be transplanted in vivo) we compared incoming and outgoing signals to the myeloid cells between the adult mouse retina, replacing the myeloid population of host upon transplantation.

The differentially enriched genes were then applied onto the three sets of CellChat objects. For example, a quantitative analysis of Apoe across the three groups (young adult, mature adult, aged) of healthy adult retina where 1) the ‘host’ myeloid is preserved, i.e. native healthy adult, 2) myeloid population is replaced with the host upon transplantation myeloid cell population, 3) myeloid population is replaced with the developing retina myeloid population.

### ssGSEA pathway analysis of integrated myeloid population

The integrated myeloid population dataset from the 5 conditions (development, adult, glaucoma model, optic nerve crush model, host upon transplantation) were used to perform ssGSEA pathway enrichment analysis using escape v. 1.10.0 ^30^ We focused our analysis on inflammation-oriented pathways from the Molecular Signature Database. Normalized enrichment score (NES) was averaged per group and pathway, and then normalized from 0 to 1 in a pathway-independent manner. The resulting table of normalized averaged NES in a pathway versus condition was visualized as polygon chart.

### WGCNA analysis of integrated myeloid population

Furthermore, the integrated myeloid dataset from the 5 conditions described above were used to perform WGCNA analysis using WGCNA v. 1.72-1 and hdWGCNA v. 0.4.03 packages. We identified modules specific to the “host upon transplantation” condition and used the top three with the highest score to build the hub genes network plot. The resulting object were saved in .rds file and these are available upon request.

### Validation of selected single-cell RNA sequencing targets by immunohistochemistry in retinal whole mounts

To validate the expression of selected differentially expressed genes identified from the analysis, immunohistochemistry analysis was performed on retinal wholemounts after RGC transplantation. Briefly, another batch of mice of the same background, age and sex were transplanted with donor RGC, and on the third day, mice were sacrificed. After enucleation, the globe was punctured with #11 scalpel at the limbus and fixed for 15 minutes in 4% paraformaldehyde at room temperature. Tissue was washed briefly with PBS to remove excess fixative, followed by removal of anterior segment and lens. Relieving cuts were made to the retina to enable flat-mounting, after which retina was extracted and washed with PBS (pH 7.4; 3 x 5 mins) and transferred to blocking solution (PBS with 10% donkey serum, 0.5% Triton-X, 1% BSA). Blocking was performed for 4 hours at room temperature, after which retinas were incubated with primary antibodies in blocking solution at 4° C for 3 days. After primary incubation, retinas were washed with PBS (3 x 15 mins), then incubated with secondary antibodies in blocking solution for 2 days at 4° C. After secondary incubation, a final series of PBS washes (3 x 15 mins) was performed, at which point retinas were flat mounted with Prolong Diamond Antifade Mountant (ThermoFisher Scientific) and cover slipped. Primary antibodies used were: Tmem119 (1:500, Synaptic Systems #400 004), Spp1/OPN (1:500, R&D Systems #AF808), and Apoe (1:500, Synaptic Systems #465 004). Secondary antibodies used were donkey anti-guinea pig 405 and 647 (Jackson Immuno Research #706-475-148 and #706-605-148), and donkey anti-goat 647 (Invitrogen #A32849). Slides were imaged on a Leica TCS SP8 confocal microscope, with optimization of brightness and contrast performed in ImageJ.

## RESULTS

### Healthy adult mouse retina atlas allows for reference-based studies of a defined class of retinal cells in young and aged mice

We sourced, analyzed, and integrated multiple publicly available single-cell RNA sequencing datasets from healthy adult mouse retina, establishing a ‘healthy adult mouse retina atlas’ which is now available here (link available upon publication). The atlas contains a total of 373,651 cells, from postnatal week 4 to 52-week-old mice **(Figure 1A, Figure S2)**. Using canonical cell markers, several major retinal cell classes were identified from our data including retinal ganglion cells (RGC), amacrine cells (AC), horizontal cells (HC), bipolar, rod, cone, Müller glia, immune/myeloid cells, retinal pigment epithelial cells (RPE), and vascular endothelial cells based on the expression of their respective known cell class-specific genes such as Pax6, Atoh7, Vim, Rlbp1, Otx2, Pde6h, Rho, Cd31, respectively. (**Figure 1A-C)**.

**Figure 1.**
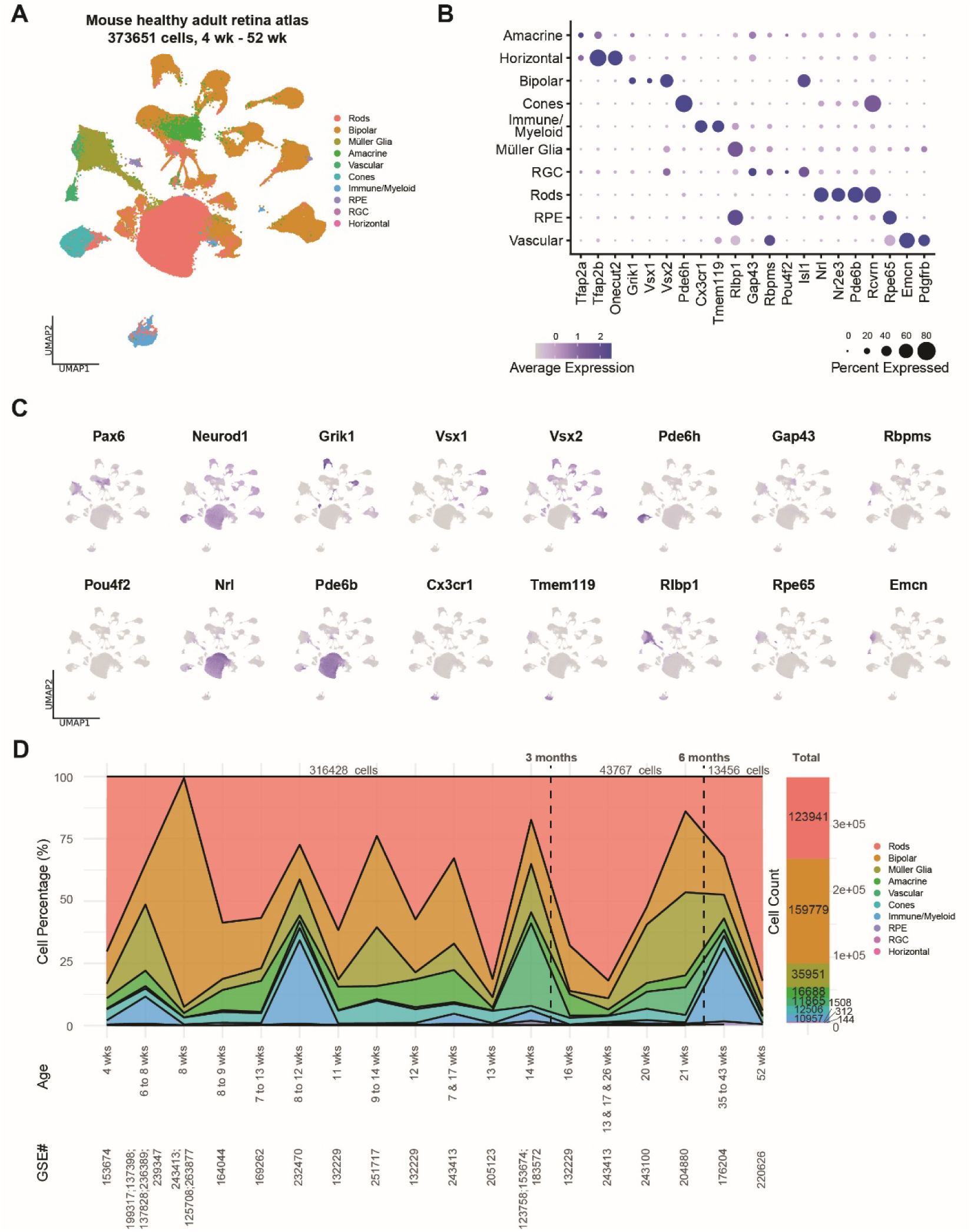
Mouse healthy adult retina atlas demonstrates high resolution of retinal cell classes over a broad timeline. **(A)** UMAP embedding of mouse healthy retina atlas (373651 cells) highlighting the major retinal cell classes identified. **(B)** Dot plot presenting the panel of canonical markers for the cell classes identified in the atlas. **(C)** Feature plots highlighting the cells expressing the class-specific markers. **(D)** Cumulative chart showing the atlas composition for the relative and absolute number of cells, timepoints distribution, and the relevant public dataset identifiers.

When the integrated data was stratified by biological timepoints (**Figure 1D**), the proportion of different cell classes varied at the different timepoints. Rods for instance constituted the largest percentage of the retinal neurons throughout the timeline. For a more streamlined downstream analysis, the data in the atlas was reclassified generating three major timeframes **(Figure S2).**

- Young adult (<=14 weeks old)
- Mature adult (15 weeks to 21 weeks old)
- Aged adult (>35 to 43 weeks and 52 weeks old)

This reclassification maintained a sufficient resolution to define different cell classes, and timepoint for in-depth subset population-oriented studies, and for cell-cell interaction analysis **(Figure S3)**. The major cell classes were maintained even after the reclassification. The Immune/Myeloid/Lymphoid cell populations contained 10,957 cells across the timeline: 9979, 205 and 773 in young adult, mature adult, and aged adult, respectively. The 10,957 of the Immune/Myeloid/Lymphoid cells represent 46 separate batches **(Figure 2A)**. Cell fate trajectory reconstruction **(Figure 2B)**, indicates two potential cell fate bifurcations, which in turn resulted in three final cell subpopulations including Dendritic cells, T cells and microglia/macrophages **(Figure 2C)**. The first bifurcation, defined by Cxcl10+ Ccl2+ Tnf+, separates immune cells and microglia/macrophages defined by Il1b+ Cybb+ Lyz2. The second bifurcation, present within the immune cell population, further subdivides into dendritic cells (Ccr7+) and T cells (Cd3e+ Cd3g+ Ccl5+ Il1b+) **(Figure 2D)**.

**Figure 2.**
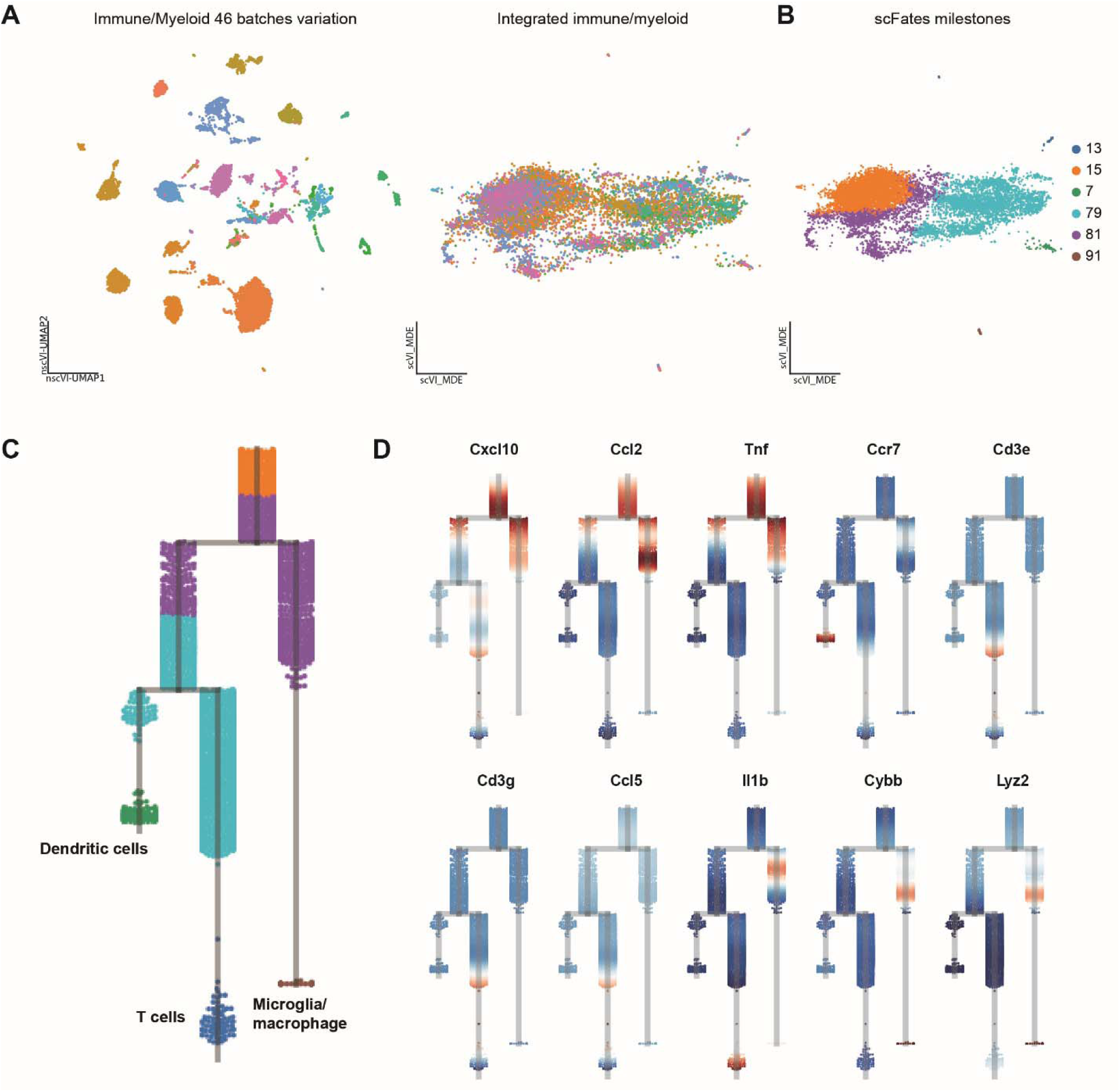
Healthy adult retina myeloid subset cell fate trajectory inference analysis decomposes the population into the major cell types. **(A)** Healthy adult retina myeloid subset (46 batches) upon scVI reembedding. The left embedding demonstrates significant batch variation, that was adjusted on the right plot of integrated myeloid. **(B)** Major milestones, or clusters, present in the myeloid subset upon scFates analysis. **(C)** Dendrogram highlighting the trajectory of milestones. Milestone 15 gives rise to two branches, with the left split into dendritic cells and T cells, while the right is true microglia/macrophage. **(D)** Dendrograms demonstrating the expression of branch-related genes, classically known for the dendritic, T, and microglia/macrophage cells.

### Host microglia respond to RGC transplantation with a continuous and reversible cell activation state

Following RGC transplantation, the host retina myeloid cell population was isolated and used for single-cell RNA sequencing (**Figure S1**). A total of 2814 cells were recovered after filtering and quality control. Automated cell annotation was performed using the previously established adult mouse retina atlas (available here: link available upon publication) as the reference atlas. Trajectory reconstruction of the annotated cells using RNA Velocity indicated a continuum of cell state upon embedding, as demonstrated by the velocity pseudo time (**Figure 3A**). In total three myeloid cell states (Homeostatic, proliferating and activated, based their expression profile) were identified i.e. the two ends referred to as the “*tip*” states (top left and bottom right ends of the trajectory) and the “*transitory*” state (the in-between the two tips), highlighted with Mellon density plot (**Figure 3A**). The trajectory demonstrates a continuous connection between the two distinct cell states of myeloid cells i.e. homeostatic and activated. We observed a bidirectional commitment of cells. The cells in the homeostatic state were capable of transitioning into an activated state, while cells in the activated state were capable of transitioning back to the homeostatic state. Importantly, these trajectories are determined by different transcriptional programs, with no major overlap on the embedding. The homeostatic and activated cell states representing the top and bottom ends of the pseudo time respectively were defined using the homeostatic and activated scores based on known genes associated with microglial ^31^ states (**Figure 3B**).

**Figure 3.**
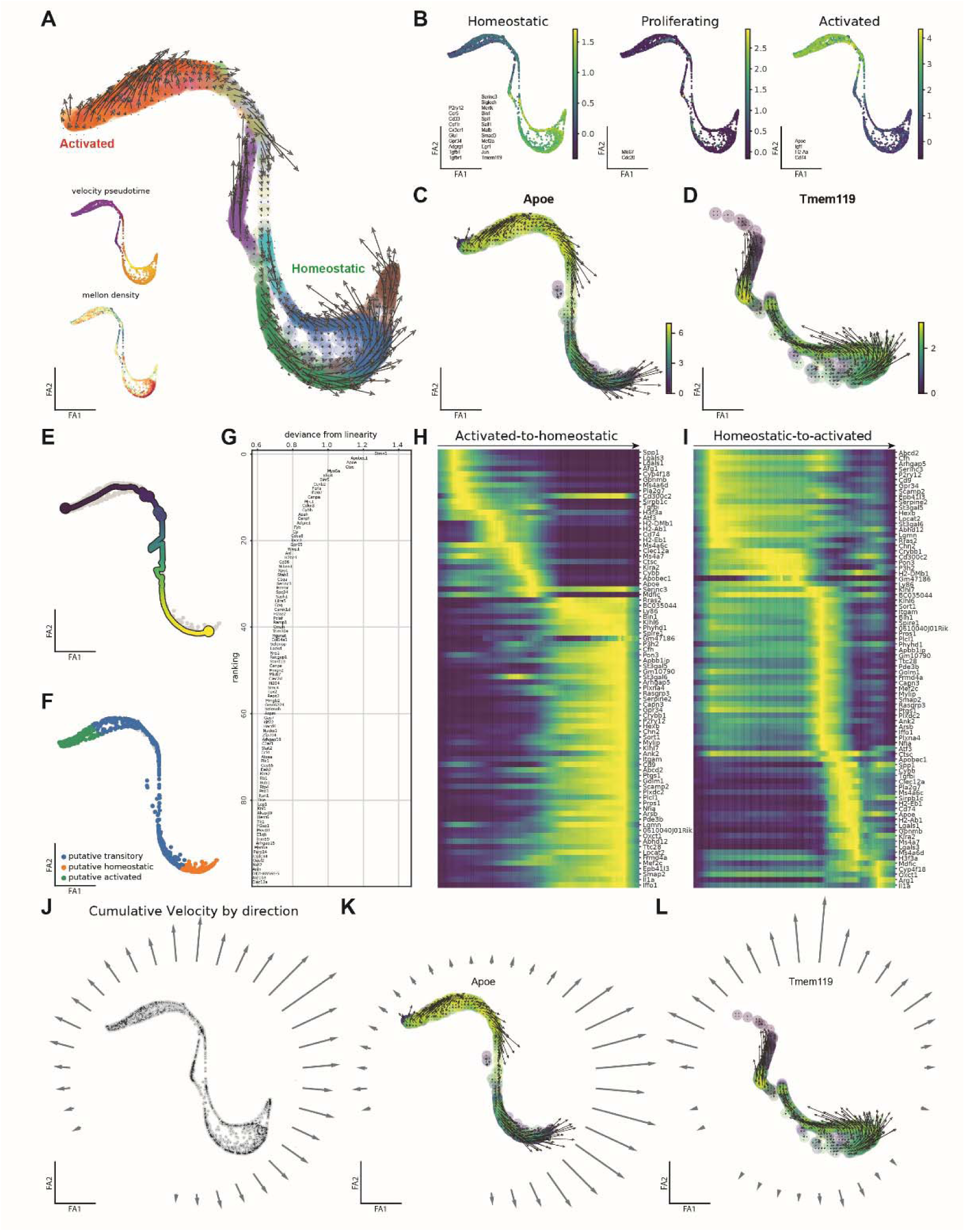
Unraveling the profiles of host microglia continuous and reversible cell activation in response to transplantation. **(A)** ForceAtlas2-based embedding with RNA Velocity vectors, demonstrating the commitment of each cell towards the homeostatic or activated state. **(B)** Featureplots presenting the cumulative scores of homeostatic, proliferating, and activated programs for the microglia/macrophage states. **(C)** Activated-to-homeostatic subset of microglia/macrophage correlates with decreasing Apoe gradient along the activated-to-homeostatic trajectory. **(D)** Homeostatic-to-activated subset of microglia/macrophage correlates with decreasing Tmem119 gradient along the homeostatic-to-activated trajectory. **(E)** Activated-to-homeostatic subset of microglia/macrophage general trajectory reconstruction. **(F)** Activation states of microglia/macrophage population identified on the subset of activated-to-homeostatic microglia/macrophage. **(G)** Variable genes identified along the activated-to-homeostatic trajectory. **(H)** Variable genes identified upon RNA Velocity analysis on the activated-to-homeostatic subset, ranked by pseudotime. **(I)** Variable genes identified upon RNA Velocity analysis on the homeostatic-to-activated subset, ranked by pseudotime. **(J)** Cumulative velocity distance distribution for the complete dataset on the ForceAtlas2-based embedding. **(K)** Homeostatic-to-activated subset cumulative velocity distances, highlighting the major direction of commitment pointing to the top left side, i.e. the position of activated microglia/macrophage. **(L)** Activated-to-homeostatic subset cumulative velocity distances, highlighting the major direction of commitment pointing to the bottom right-right side, i.e. the position of homeostatic microglia/macrophage.

Using *Tmem119* ^32^ and *Apoe* ^9^ genes as markers of homeostatic and activated microglia respectively, we show that activated-to-homeostatic transitioning (**Figure 3C**) is defined by a decreasing expression of *Apoe*, while the transitioning from homeostatic-to-activated (**Figure 3D**) is defined by a decreasing expression of *Tmem119*. Activated-to-homeostatic subset was then used to build a common supervised trajectory starting from the most activated cell to the least activated, identifying three cell states i.e. ‘putative transitory’, ‘putative homeostatic’ and ‘putative activated’ (**Figure 3E, F**). The trajectory displayed with the gene IDs as a deviance from the healthy, with the genes located at the top of the trajectory defining the activated cell state, and the genes at the bottom defining the homeostatic cell state (**Figure 3G**). In the same manner but for the RNA Velocity trajectory, with the idea of including potential local perturbations, we profile the activated-to-homeostatic trajectory (**Figure 3C, H**), and the homeostatic-to-activated (**Figure 3D, I**) trajectory with the variable genes along them. Transitioning from the activated-to-homeostatic (**Figure 3H**) was defined by the suppression of genes associated with activated microglia such as *Lgals3*, *Spp1*, *Cybb*, and *Apoe*, while transitioning from the homeostatic-to-activated state was defined by a suppression of a microglia homeostatic gene *P2ry12*, and at the same time upregulation of *Apoe*, a gene associated with microglia activation (**Figure 3I**).

For a quantitative assessment of cell commitment towards either the activate (preCS_a_) or the homeostatic (preCS_h_) state, and to profile the homeostasis balance between the two states, we introduced a numerical *“commitment score*” (CS) which summarizes RNA Velocity vectors that follow a given direction either towards homeostatic or activated states. The cumulative velocity scores representing the distribution of vectors for cell commitment for the homeostatic, and for the activated microglia are shown in **Figure 3J**. They were further separated into two subsets: homeostatic-to-activated and activated-to-homeostatic, and for both trajectories, the preCS_a_ and preCS_h_ scores were quantified (**Figure 3K, L**).

We explore the state commitment of myeloid population (CS) as a ratio of the homeostatic and activated commitment scores. The value of unnormalized commitment score (unCS) = 9.335. We normalize this value by assessing the number of cells within each state. The value of normalized commitment score (nCS) = 8.066, meaning the general trend of the population commitment is highly shifted towards the homeostatic state. The commitment of cells towards the homeostatic state is between 8.066 and 9.335 times higher compared to the commitment towards the activated state.

### Differential expression of selected genes in the host retina following xenotransplantation of RGC

The expression of genes identified from the analysis of our single-cell RNA-seq data was validated by immunohistochemistry. Retinal whole mounts were stained for Spp1, Tmem119 and Apoe three days after RGC transplantation **(Figure 4A-R)**. Close to the injection site, we noted a localized increase in density, and a change in morphology of the host GFP+ myeloid cells. These cells were noted to be less ramified and more ameboid in shape compared to the same cell type further away from the injection site. Expression of the homeostatic microglial marker Tmem119 was absent at the injection site but was present in the ramified microglia elsewhere in the tissue. The DAM marker Spp1 was expressed in GFP+ myeloid cells at the injection site (B, F) but not elsewhere (G, K) (of note, Spp1 is also expressed by a subpopulation of retinal ganglion cells). Additional immunohistochemical investigation (L – R), both at the surface (M – O) and within the retina (P – R) reveled localized increase in Apoe expression at the injection site. These findings confirm that host myeloid cells in proximity of transplanted RGC change their morphology and upregulate classic DAM markers Spp1 and Apoe.

**Figure 4.**
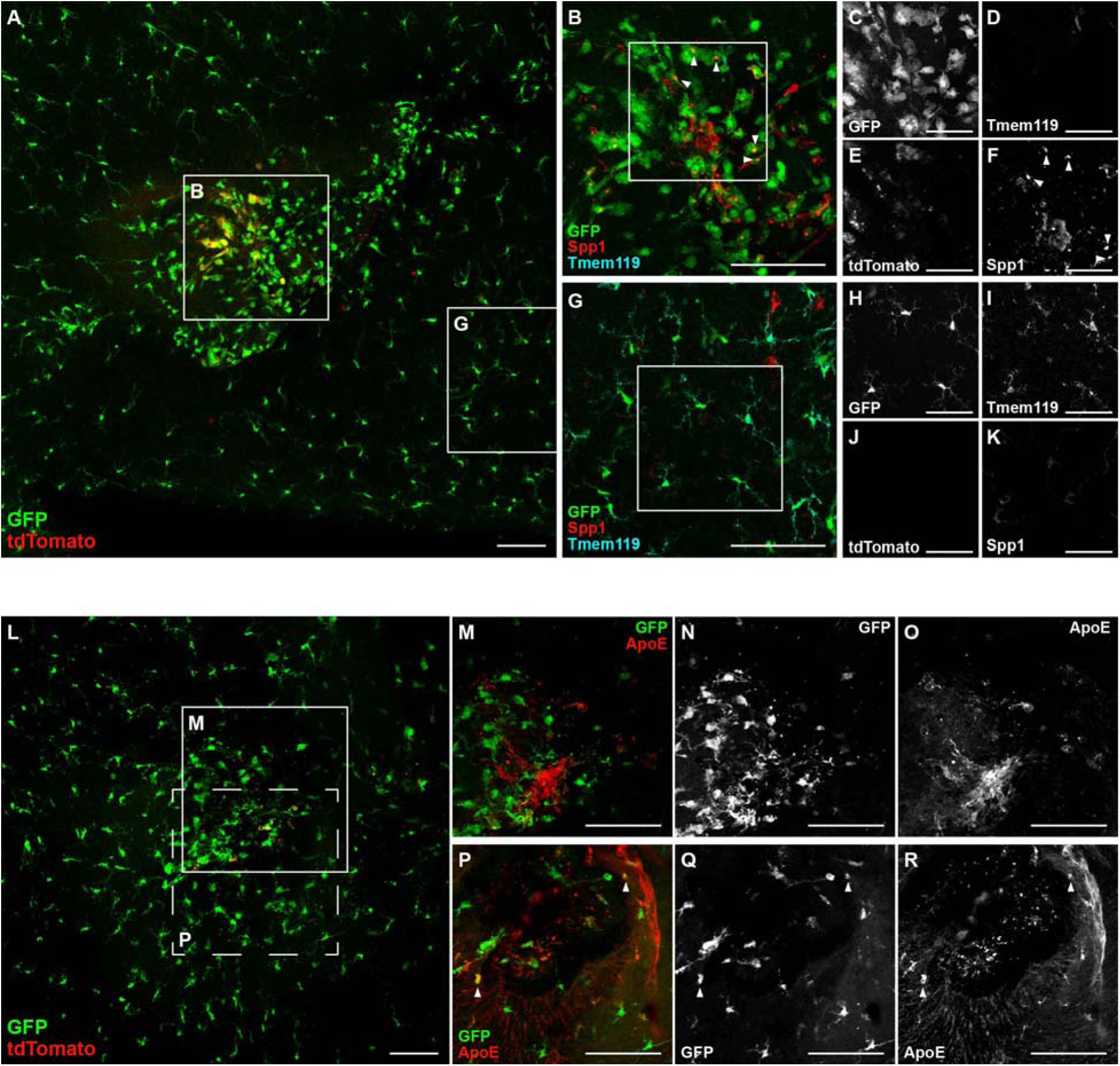
Injection of tdTomato^+^ hESC-derived RGC results in a localized increase in density of Cx3cr1-GFP^+^ myeloid cells (microglia and macrophages), and loss of characteristic homeostatic microglia traits including ramified morphology and expression of the marker Tmem119. **(A)** Injection sites, identified by tdTomato **(A, E)**, display highly localized increases in density of GFP-labeled myeloid cells **(B)**, along with transition to amoeboid morphology **(B, C)**, in contrast to nearby regions of the retina **(G, H)**. Likewise, expression of homeostatic microglial marker Tmem119 is absent within the injection site **(D)** but persists in ramified microglia elsewhere in the retina **(I)**, whereas the DAM marker Spp1 is found in GFP+ myeloid cells at the injection site **(B, F)** but not those elsewhere **(G, K)**. Additional immunohistochemical investigation **(L – R)**, both at the surface **(M – O)** and within the retina **(P – R)** reveals localized increase in Apoe expression at injection sites, including in amoeboid GFP+ myeloid cells below the retinal surface **(P – R)**. Scale bars = 100 µm **(A, B, G, L – R)**, 50 µm **(C – F, H – K)**.

### *In silico* integration of myeloid/lymphoid cell populations from the developing, adult, and damaged mouse retina reveals novel mechanisms of host response to RGC transplantation

We integrated myeloid cell populations (**Figure 5A**) from the following conditions:

a. Developing retina available here: link available upon publication
b. Healthy adult retina available here: link available upon publication
c. Glaucoma model (microbead-induced IOP elevation) available here: link available upon publication
d. Optic nerve crush model available here: link available upon publication
e. Myeloid cell population from the host retina after transplantation of human RGC

**Figure 5.**
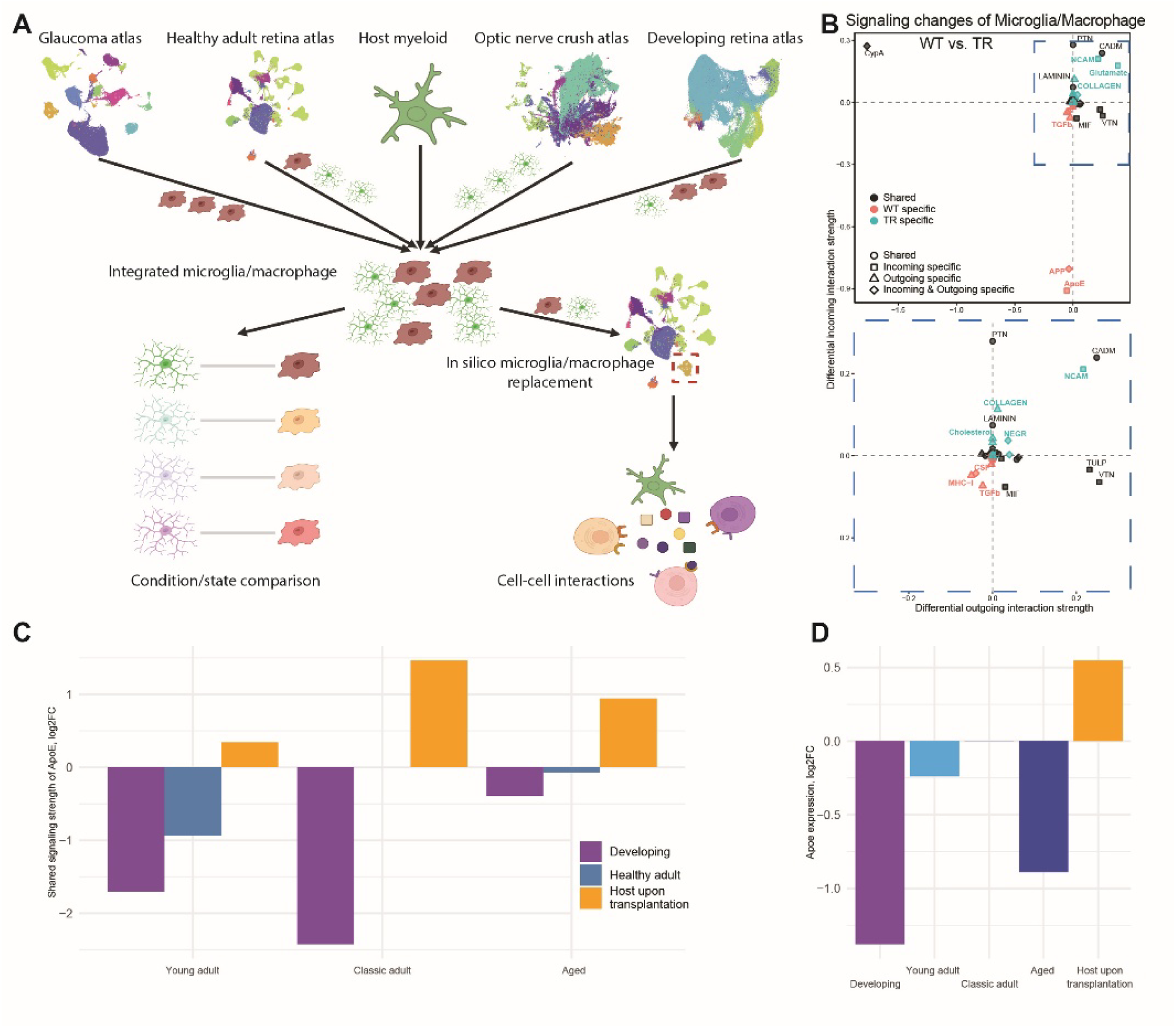
In silico integration of the developing, adult, and damage conditions myeloid population reveals the mechanism of host response to transplantation. **(A)** Abstract sketch demonstrating the in silico pipeline for the integrated microglia/macrophage downstream analysis. **(B)** Cell-cell interactions comparison of healthy adult mouse retina microglia/macrophage versus host upon transplantation microglia/macrophage in the 20 weeks old healthy adult mouse retina surrounding. **(C)** Plot representing summarized incoming and outgoing signaling strength of Apoe signaling in developing, healthy adult, and host upon transplantation microglia/macrophage in the surrounding of young adult, mature adult, and aged retinal cell classes of mouse healthy adult retina atlas. **(D)** Apoe expression in microglia/macrophage of developing, young adult, mature adult, aged, and host upon transplantation conditions.

*In silico* experiments were performed by artificially replacing a given myeloid cell population of one condition with those from another condition, for example, moving myeloid cell population from the optic nerve crush condition, and placing them into the adult healthy retina. This *in-silco* experimental approach allows us to create a unique biological model for exploring cell-cell interactions between different conditions.

For the *in silico* experiments, we used the “20-week timepoint” dataset to mimic cells from an adult mouse host retina, one into which RGC would typically be transplanted. We compared signaling cascade of myeloid cells between the native healthy adult 20-week timepoint, with the same condition but with the native myeloid cell population replaced by the myeloid cells from the host. Cell-cell interactions analysis revealed several unique signaling mechanisms including increased signaling strength in NCAM, Glutamate, and COLLAGEN, and decreased signaling strength of TGFb, Apoe, and APP (**Figure 5B, C**).

We also profiled cell-cell interactions in three additional groups including:

- Young adult (<=14 weeks old)
- Classic adult (15 weeks to 21 weeks old)
- Aged adult (>35 to 43 weeks and 52 weeks old)

And used three sets of myeloid cell populations to populate the healthy adult atlas i.e.

1. Native healthy adult myeloid
2. Developing (E10-P8),
3. Host myeloid from the retina upon human RGC transplantation.

Apoe was found to be differentially expressed in microglia/myeloid across the conditions. Apoe was downregulated in development, in the young adult, and in the healthy aged animals, and upregulated in the host upon RGC transplantation (**Figure 5D**). This Apoe differential expression pattern is in line with reported expression of Apoe in DAMs ^10^. WE observed the lowest Apoe level in the developing myeloid cells and the highest in host-upon-transplantation condition. We also observe continuous increase with aging win Apoe expression increasing in young adult, then aged adult, and highest in the mature adult. These findings indicate that the developing and young adult retina may be the most amenable for the transplantation, as retinal cells at this age showed the least activation response to transplantation. With the observed Apoe expression pattern across the different conditions, we then correlated Apoe signaling strength in the myeloid cell population with the Apoe expression between the conditions of developing, healthy adult (young, mature, and aged), and host upon transplantation myeloid populations (**Figure S4**).

### Myeloid cell populations across the different conditions have a differential microglia activation profile

To evaluate microglia activation, inflammatory response, and other pathways we perform Single-Sample Gene Set Enrichment Analysis (ssGSEA) pathway analysis from the following conditions of the myeloid population: developing (E10-P8), healthy adult (4-52 wk), host upon transplantation (6 months), optic nerve crush <= 4 days (12 hours, 1 day, 2 days, 3 days, 4 days), optic nerve crush > 4 days (7 days, 14 days), and microbead glaucoma model (7-22 wk) (**Figure 6A**). The analysis focused on inflammatory pathways, and showed that microglia/macrophages from the developmental cell state had the lowest normalized enrichment score (NES) in each of the analyzed pathways. The highest NES was observed in the host upon transplantation condition for most pathways.

**Figure 6.**
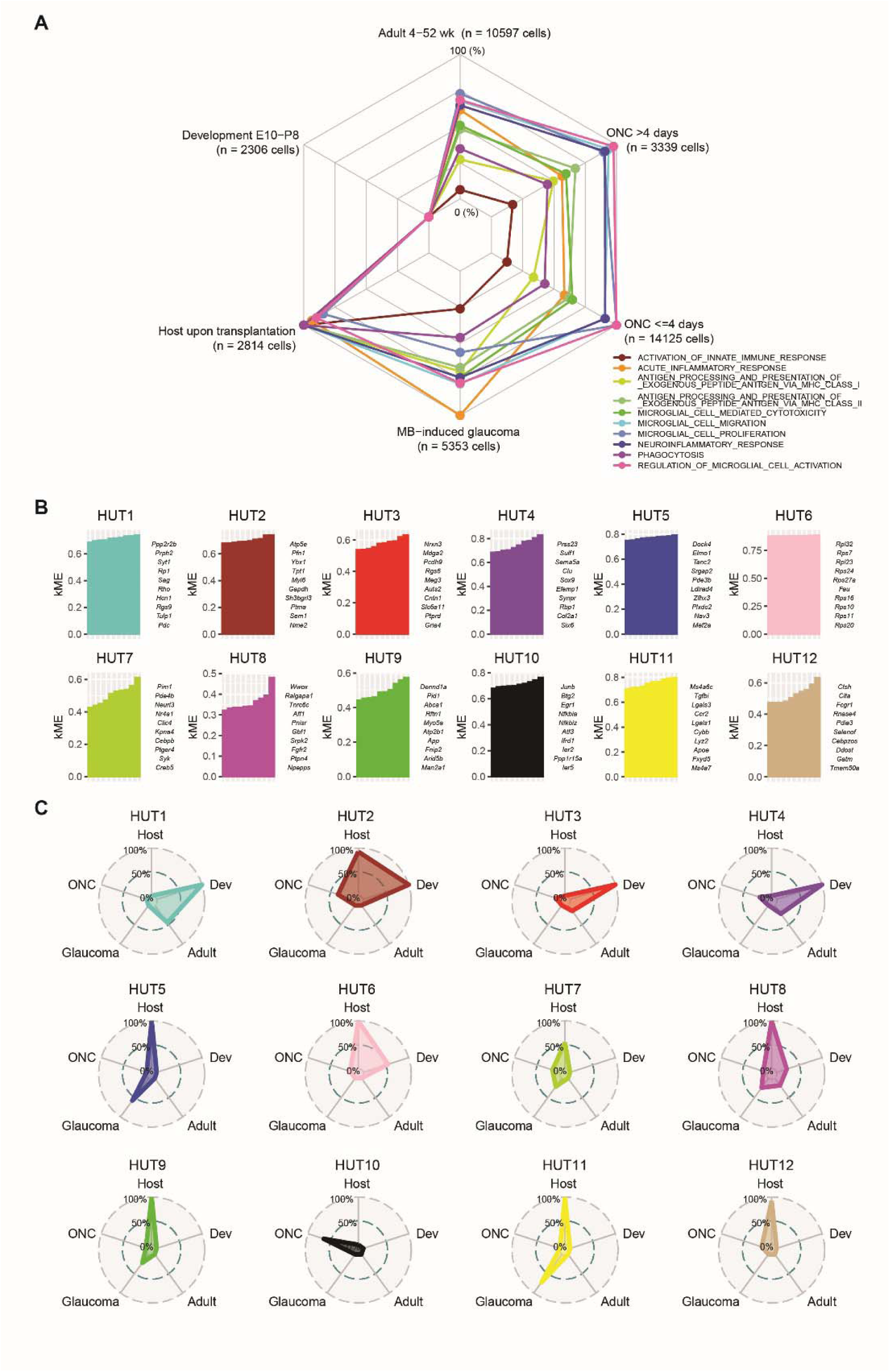
Integrated myeloid dataset comparison demonstrates differences in microglia activation. **(A)** Polygon chart representing the results of ssGSEA pathways analysis on inflammatory-oriented pathways in the developing, healthy adult, optic nerve crush, glaucoma model, and host upon transplantation microglia/macrophage conditions. **(B)** WGCNA analysis represented in 12 modules with top 10 hub genes in each module. **(C)** Polygon chart demonstrating the modules distribution across the conditions from the integrated myeloid dataset.

Of the pathways analyzed, GOBP_ACUTE_INFLAMMATORY_RESPONSE had the highest NES in the glaucoma condition, while GOBP_REGULATION_OF_MICROGLIAL_CELL_ACTIVATION along with GOBP_MICROGLIAL_CELL_PROLIFERATION had the highest NES for the ONC condition. This enrichment indicates a similarity in microglia activation pattern between disease conditions and host cell response to RGC transplantation (**Figure 6A**).

A closer look at the genes involved in the enriched pathways indicated a differential expression pattern. In GOBP_ACTIVATION_OF_IMMUNE_CELLS pathway, several genes were differentially upregulated in the host including: Sting1, Arrb2, Nlrp6, NfkbiL1, Mark4, Akt1, Myd88, TLr8, and Sec1411 **Figure S8**. These genes were uniquely upregulated in the host retina post transplantation but were suppressed in the MB-induced glaucoma model, in the ONC model, and during development. In this manner, we provide the complete list of the genes along with their relative expression involved in the pathway analysis between the conditions, in **Figure S8**.

Using the same dataset we performed whole genome correlation network analysis (WGCNA) in an unsupervised manner. The analysis revealed twelve HUT modules (**Figure 6B-C**, **Figure S5A, Figure S7**), with HUT1 being specific for the healthy (developing and adult), HUT3 and HUT4 for developing, HUT6, HUT8, HUT9 and HUT12 for host upon transplantation, HUT5 and HUT10 for optic nerve crush condition, and HUT11 for host upon transplantation and MB-induced glaucoma.

HUT11 represents an activation module, with several of the hub genes including Ccr2, a receptor for Ccl2 a chemoattractant important for macrophage and monocyte recruitment, Lgals1, Cybb, Lyz2 and Apoe, genes associated with microglia activation in disease (**Figure 6C, Figure S6**). This module was significantly expressed in host upon transplantation and glaucoma conditions **(Figure S5)**. The optic nerve crush condition was represented by HUT10 and it includes genes as Junb, Egr1, and Atf3, known transcription factors that modulate activity of several other genes. The healthy condition (developing and adult) was characterized by HUT1, which included several genes including *Rp1*, *Sag*, *Rho*, *Tulp1*, and *Pdc*, which genes are implicated retinal developmental diseases such as retinitis pigmentosa. From the selected the modules with the highest score in host upon transplantation condition (**Figure 6C**) we build the hub gene network (**Figure S5B**), where HUT2, HUT6 and HUT11 form a distinct cluster, and genes from HUT8 and HUT9 form another cluster.

## DISCUSSION

The success of RGC restoration by transplantation depends on several factors including the state of the host microenvironment. In this study, we aimed to understand mechanism regulating host myeloid cell response following RGC transplantation. We utilized a combination of single-cell RNA sequencing to profile the myeloid cell population, advanced integrated transcriptomic analysis, and immunohistochemistry to characterize the host cell reactivity/morphology after RGC transplantation.

Single-cell datasets of the retina have been integral for analyzing single-cell RNA sequencing data, however, they face challenges such as the lack of representation of rare cell types, not being representative enough of the different experimental conditions in health and in disease, and the need for validation of the data.^33,34^ For a comprehensive understanding of developmental, and of disease processes, a large-scale atlas integration approach with high resolution at the single cel level is paramount. This can be accomplished by collecting multiple datasets that will provide the necessary coverage in terms of different cell identities and cell-states in each of the individual studies. This labor-intensive approach can simply be overridden by utilizing a well curated, indexed reference atlas which allows for the integration of the new “experimental” datasets into the existing atlas. A major requirement for such a comprehensive reference atlas is that it should contain all the different cell states and cell identities represented in the given study dataset. In the context of the present study, a useful atlas should cover all myeloid cell classes and different cell states including homeostatic and activated states. As such, to facilitate a better analysis of our complex dataset, we have established a well curated and representative adult mouse retina single-cell atlas using publicly available single-cell RNA-seq datasets. We generated an atlas containing a total of 373,651 cells, containing samples from mice ranging from 4 to 52 weeks of age. To the best of our knowledge, this adult mouse retina atlas is the largest of its kind at the time of this writing, covering a range of cell types and experimental conditions. This atlas can serve as a reference for interrogation functional studies involving the retina and optic nerve (aging, in-depth cell class and type analysis, regeneration potential). Furthermore, the atlas can be used as a control dataset for multi conditional studies, and/or serve as a reference tool when analyzing other single-cell RNA-seq datasets e.g. for automated cell annotation.

We were able to successfully assemble the scRNAseq atlas and integrate and normalize datasets, overriding the batch effect, arising from different chemistry, platforms, sequencing approaches and experimental designs. One should caution when using such Atlas: it allows to study the gene expression and trajectory reconstruction when built-in controls are used - mitochondrial, ribosomal, or MALAT1.^35^ However, such an atlas should not be used to frequency-related analyses, lifespan, aging, and other temporal-oriented questions. Nevertheless, large-scale public data integration into the atlas format, with multiple metadata variables (age, gender, race, ethnicity, supporting disease) creates a wide field for in silico testing of the hypotheses, especially population- and clinical-oriented. This atlas could also serve as a control, “natural history” group for multiple potential studies, without the requirement of collecting the control samples in an independent experiment.

Furthermore, this mouse control adult retina atlas, spanning 4 to 52 weeks of age allows to study the maturation and aging of individual cell classes. We demonstrated a continuity of timepoints in the integrated atlas, and this can enable cell class-oriented studies: for instance, a user may subset the amacrine cell class and explore the transcriptional changes occurring over time. The metadata such as gender, sequencing depth and platform, and multiple other variables, are important for both biologically- and technically oriented studies. One example is a sample variance analysis ^36^, which can be now be performed within the atlas.

Establishment of the reference atlas was then followed by annotating our single-cell RNA seq data, with reference to the atlas. For this study, we use the terminology “myeloid” to describe all possible myeloid cell populations collected from the retina before and after donor RGC transplantation, and these include Cx3cr1+ cells (microglia, resident macrophages, monocyte-derived macrophages), Cx3cr1^low^ activated cells with an activated transcriptional profile, and other myeloid cells. Due to the complexity of microglia and macrophage profiling, we use the term “Microglia/macrophage” to highlight the population of our interest in the analysis. Since this study is function-oriented and activation-oriented, we did not perform further separation to microglia and macrophages when it comes to cell fate trajectory inference or any other downstream analysis.

Upon cell trajectory analysis we confirm an existing paradigm that microglia/macrophage population of the retina is a dynamic system, and each cell within this system may exist in either the “homeostatic”, “activated” or the “transitory” state. We also profile the continuity and bidirectionality of the changes. These confirmed features align with the previous discoveries performed on microglia in healthy wild type mouse retina and the retina of microbead-induced glaucoma mouse model, ^37^ proposing the activation mechanisms to be flexible, reversible and bi-directional. Previously demonstrated cell fate trajectory reconstruction methods highlight the system to be controllable, and the homeostatic-activated homeostasis to be a unique condition-related feature. The application of cell fate trajectory inference methods ^38^ assumes that we are working with the dynamic system. Here, we applied RNA Velocity method to reconstruct the evolution of host microglia/macrophage population with the dynamics in the activation profile in response to RGC transplantation. The myeloid population of the healthy adult retina can also be analyzed with similar strategy due to the ongoing myeloid cell renewal throughout life. This creates necessary dynamic, that permits potency-based deconvolution and allows to apply scFates algorithm.

The decision to use unnormalized or normalized commitment score should be based on an experimental question and design. Since the main difference between these two metrics is in the cell number of adjustment, it is important to know the reason of such cell number variation. This could be the true condition-related frequency difference, sample variation, or the library preparation-related issue. Unless it is possible to determine the reasoning, the true commitment score value lays between the unnormalized and normalized, where stronger condition-related frequency change is a hint to look closer to the normalized commitment score value, and stronger batch-related frequency change can be more associated with the unnormalized value. In the real case scenario, additional methods to identify the reasoning of frequency change, such as Bayesian models application for the dataset composition analysis, could help in finding out the real commitment score.

We observe a well-defined transitory state of microglia/macrophages between the homeostatic and activated tips. Our analysis suggests the presence of two transitory populations, representing the directionality of these changes: homeostatic-to-activated and activated-to-homeostatic. While these two populations are very similar in their transcriptional profile, the Euclidian distance suggests that these transitory programs are different and are not the same program executed in the opposite direction. In our experimental design with the focus on microglia/macrophage activation, we expect the bi-directional nature of changes. However, we appreciate the several discovered populations that include pro-angiogenic and IFN-response microglia/macrophages. Yet, we observe these populations to be blended with the rest of the microglia/macrophage population, suggesting that microglia/macrophages potential to activation is universal and function-agnostic. Further, high-resolution studies could provide an answer to the question of bi-directional cell state transition.

Through unsupervised clustering not based on known microglia-specific genes, we confirm previous observation that *Tmem119* and *P2ry12* are markers of homeostatic microglia. ^39,40^ The activated state was characterized by upregulation of *Apoe*, *Spp1*, *Lgals3*, *Lgals1*, *Cybb*, *Atf3*, *Cd74*, *Il1a*, and *Apobec1* and downregulation of *Tmem119* and *P2ry12*. Interestingly, this activated expression pattern is highly similar to that of disease associated microglia (DAM), also known as neurodegeneration-associated microglia (MGnD) which were first described in the context of Alzheimer’s disease, multiple sclerosis, and amyotrophic lateral sclerosis. ^19,41^ The same transcriptional signature was subsequently described in the retina in glaucoma and photoreceptor degeneration mouse models, as well as human glaucoma and age-related macular degeneration retinal tissues. ^9,42^ Interestingly, these prior studies have demonstrated that the DAM/MGnD signature can be induced by microglial phagocytosis of apoptotic or damaged neurons *in vivo* in the brain and in the retina.^19^ Furthermore, the same DAM-like signature has been identified in the developing murine retina, where it is thought to result from the developmentally appropriate microglial elimination of excess apoptotic RGC. ^43–46^ Thus, it seems likely that the DAM-like signature seen following human RGC transplantation results from retinal microglia phagocytosing transplanted RGC, and that modulating this myeloid cell response could enhance transplanted RGC survival.

Cell-cell interaction analysis with CellChat, applied in the manuscript, should be performed carefully, when applying it on the multibatch data. Due to the different composition of datasets (batches), it is important to adjust for the population presence and its size. In our case, we did not include the timepoints that were missing any of the major retinal classes, since it affects the general pool of ligands and receptors and may pull the picture to the false results. We have also excluded the timepoints with low abundance of microglia/macrophage cells. Nevertheless, even these filters do not promise the real cell-cell interactions picture. Particular timepoints may be outlying due to the different platforms, sequencing depth or coverage in the study. In this case, averaging or binning the data may be a solution to override the noise.

The GSEA pathway and WGCNA analyses allowed us to quantitatively compare several signaling pathways related to inflammation and their enrichment scores across multiple conditions. Our findings suggest that human RGC transplantation is a stronger stress factor compared to optic nerve crush or microbead-induced glaucoma and is the strongest across all conditions. Upon transplantation, microglia/macrophages upregulate MHC-I and MHC-II receptors. Their acute inflammatory response is very similar as in glaucoma model condition, yet different compared to optic nerve crush model. Also, transplantation is the only condition among those studied that resulted in microglia/macrophages to extremely increase phagocytosis function. However, we also confirm microglia activation in optic nerve crush and glaucoma model conditions, that could be one of the mechanisms of RGC death. In contrast, the behavior of developing microglia/macrophages is completely different from any other condition studied. Unlike healthy adult, optic nerve crush model, glaucoma model, or post-transplantation condition, developing microglia/macrophages have the lowest inflammatory potential.

Of the genes involved in GSEA pathway analysis, stimulator of interferon response cGAMP interactor 1(Sting1) and Arrestin beta 2 (Arrb2) are of particular interest. Sting1 is expressed by several cell types of the retina including retina pigment epithelial cells (RPE), photoreceptors, Müller glia and in microglia, and signaling of this pathway is important in health and in disease. In disease, Sting1 signaling is implicated in several diseases including diabetic retinopathy, and age-related macular degeneration where it is thought to induce senescence in endothelial cells leading to disease progression. ^47^

^48^ In animal models of glaucoma, studies have shown that targeting cGAS-STING pathway is neuroprotective. Zhang et al., showed that STING is activated in damaged retinal ganglion cells, and is associated with increased inflammation, DNA damage, and mitochondrial dysfunction. ^49^ In relation to the observed Sting 1 expression in our study, work by Liu et al., showed that Sting 1 is markedly activated in retinal microglia, and that pharmacological targeting cGAS–STING–IFN-I is neuroprotective. ^48^ In a murine model of allogeneic transplantation, Bader et al., showed that Sting is plays an important role in the regulation of of murine graft-versus-host after allogeneic hematopoietic stem cell transplantation. Studies like these indicate the important role of Sting signaling in transplantation, and as such warrant further investigation in the context of RGC transplantation. ^50^

Arrb2, another gene we found to be uniquely upregulated in the host post transplantationbelongs to the arrestin protein family, reported to dampen cellular responses to a range of stimuli, specifically via their G-protein-coupled receptors. Arr2 is thought to modulate several functions in health and in disease. In disease, Arrb2 is shown to promote inflammation in skin allergy^51^, and in Alzheimer’s disease, Arrb1 and Arrb2 expression was found elevated in AD patients.^52^ Using an HSV-1 model, Zeng et al., showed that Arrb2 inhibits activation of NF-kB signaling in microglia, favoring an ‘M2’ microglia phenotype, attenuating inflammation.^53^ In our model, it is likely that upregulation of Arrb2 is in line with the reports of its role in attenuating inflammation, but this remains to be tested.

Other genes of interest uniquely upregulated in the host post transplantation include Myeloid differentiation primary response protein 88 (MyD88), a well-defined component of the innate immune system, with a significant role in TLR/IL-IR-mediated signaling. ^54^

It is widely accepted that local inflammation greatly impacts the regenerative ability of neurons following acute injury. Using a zebrafish regenerative model for example, Rumford et al., showed that MyD88 upregulation in microglia negatively impacts cell survival, and differentiation following acute retinal damage.^55^ It is likely that upregulation of MyD88 in the host negatively impacts survival of the transplanted RGC. These findings show that MyD88 warrants further investigation as a target to promote neuronal survival post trans plantation.

In this manuscript, we also introduce a concept of the “commitment score”. For the microglia/macrophage population it is calculated based on the two-directional commitment model: the cells studied can go either to the activated or to the homeostatic state. However, this paradigm can be expanded further to quantitatively describe multi-directional systems. For example, mouse and human PAX6+ retinal progenitors give rise to amacrine, horizontal, and retinal ganglion cells. ^56^ ^57^ ^58^ Other developing systems including neural crest, brain, PBMC may contain three and more final cell states arising from the same progenitor state. In this case, a solution would be to build the force-directed layout, such as ForceAtlas2, or applying the circular projection with the number of vertices equal to the number of final states ^59,60^ With that, each sector of the plot will have equal weight, when quantifying the cumulative velocity distance and measuring the commitment score.

Our analysis identifies the most accepting host environment for donor neurons transplantation in the settings of host microglia activation. Our observations from developing microglia and host environment correlate with the literature findings showing improved survival and extensive integration of donor neurons when transplanted into the developing retina. Thus, for multiple inherited disease, including retinitis pigmentosa, Leber congenital amaurosis (LCA), Stargardt disease, and cone-rod dystrophies, transplantation in the developing system settings, could be a promising solution in absence of other therapy. Additionally, transplantation in the developing system settings allows further fundamental research: while the settings allow longer lifespan of donor neurites, they may reveal their full functional potential, if any, without the risk of being arrested due to the transplantation/neurogenesis in regeneration-unfriendly host environment. ^4,61–67^

Overall, our work underlines the importance of modulating host innate immunity, including microglia and macrophages, to improve neuron transplantation into the retina. The advanced integrated transcriptomics analysis showed that retinal microglia activation in response to neuron transplantation is profound and more dramatic than microglial response in RGC damage models. However, we also demonstrate the reversibility of this process using the pseudotime analysis, suggesting a molecular roadmap to decrease the activation of host myeloid population and increase the return to the homeostatic state.

## Data availability statement

The host upon transplantation single-cell RNA sequencing data generated in this study is deposited and available on GEO under the GSE285564 reference number.

The code generated in this study was deposited on GitHub, and the generated mouse healthy adult retina atlas is available on CELLxGENE. The links for the deposited code and data will be available upon publication and currently are available upon request. Researchers interested in contributing data for future versions of the reference atlases should contact the corresponding authors directly.

## Author contribution

**E.K;** Bioinformatics analysis, experimental design and lead first author. **A.M**; Tissue dissociation, single-cell suspension preparation, cell sorting and sorted population characterization; library preparation and logistics for sequencing. sample processing and second first author. **N.R**; - Cell differentiation; **V.V.M;** - RGC transplantation and tissue dissection, histology. **E.K**, **S.V**, **N.B**, **E.L** – Collection, parsing and processing of public single-cell RNA sequencing datasets; **P.F.C**, and **G.B** – Immunohistochemistry and imaging; **M.A.M** and **P.B** conceived the study. **E.K**, **A.M**, **M.A.M** and **P.B** wrote the manuscript. All authors approved the final manuscript.

## Supplementary material

**Figure S1.**
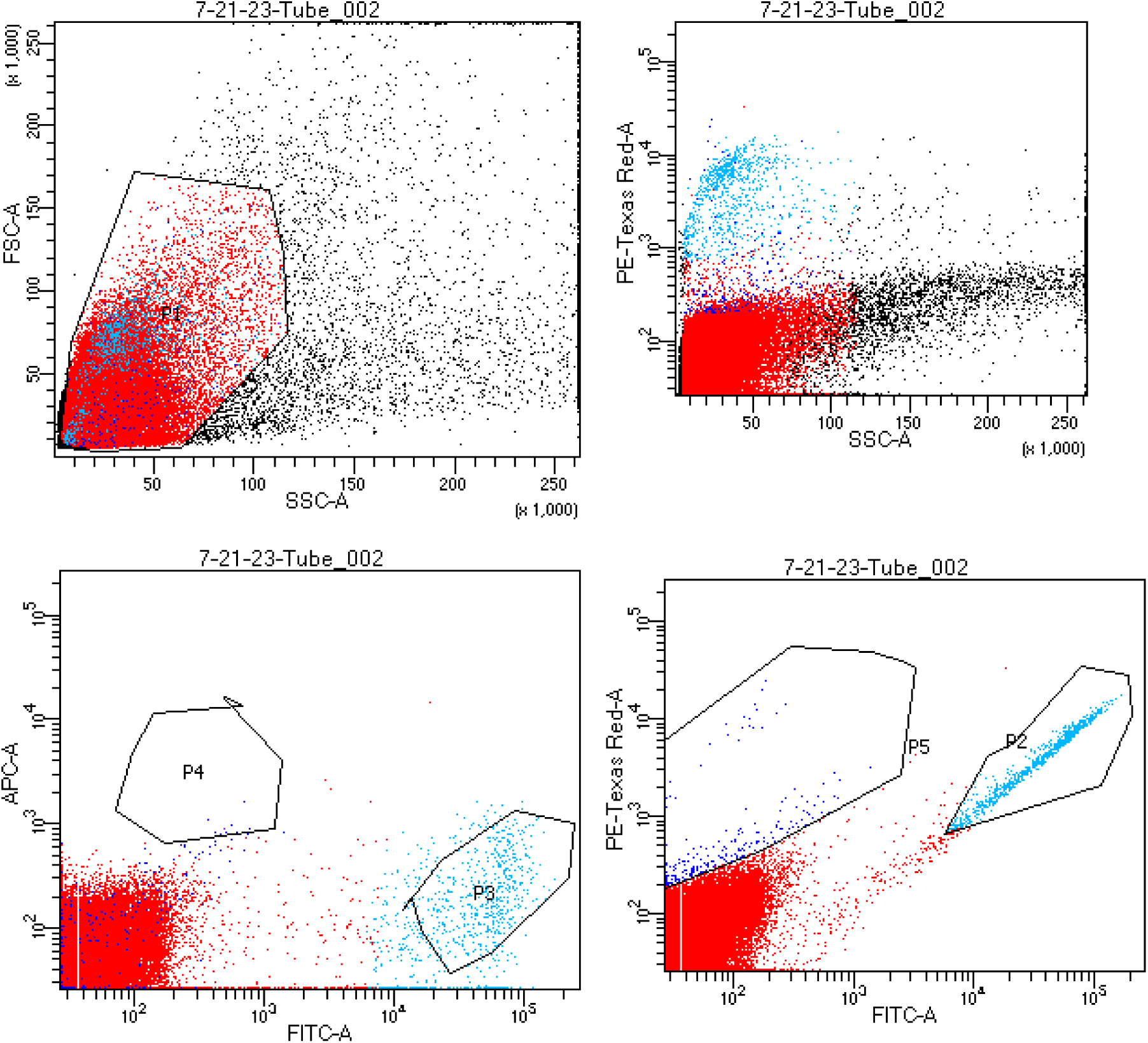
Sorting RGC (TdTomato positive) and Microglia (GFP positive) cells using FACS.

**Figure S2.**
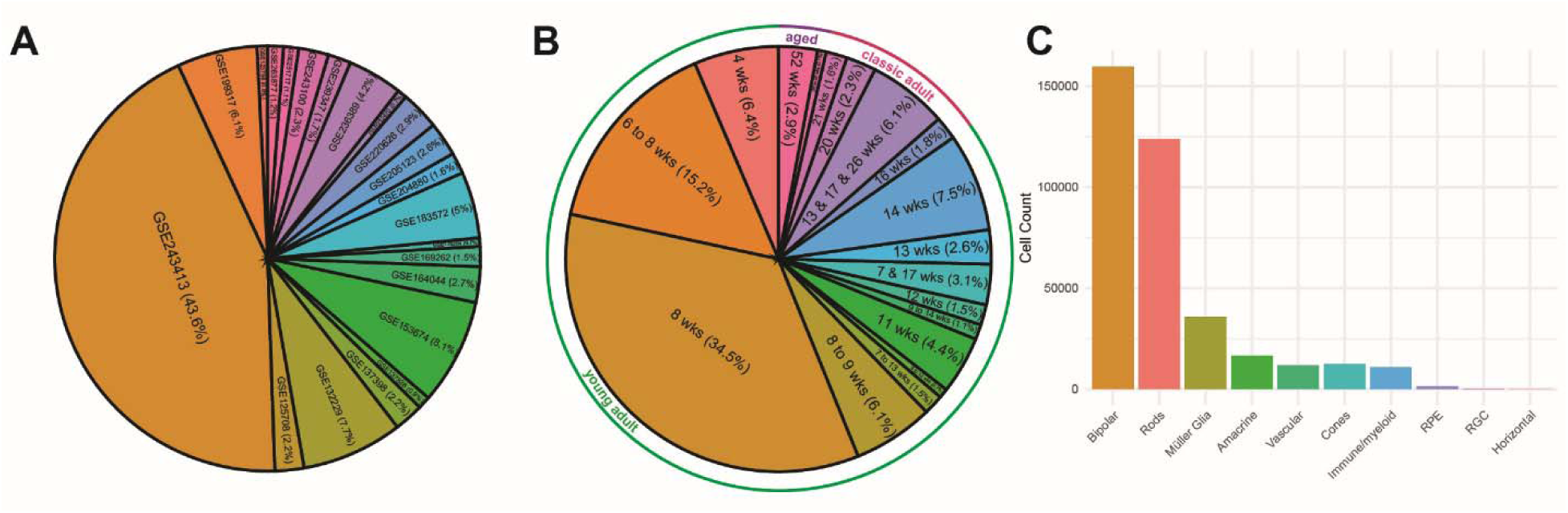
Metadata composition of mouse healthy adult retina atlas. **(A)** Pie chart demonstrating the public datasets contribution into the atlas. **(B**) Pie chart representing the timeline and timepoints abundance of the atlas. Timepoints are separated into three functional groups: young adult (4 to 14 weeks), mature adult (16 to 21 weeks), aged (rest up to 52 weeks). **(C)** Barplot highlighting the cell classes abundance in the atlas in the absolute values.

**Figure S3.**
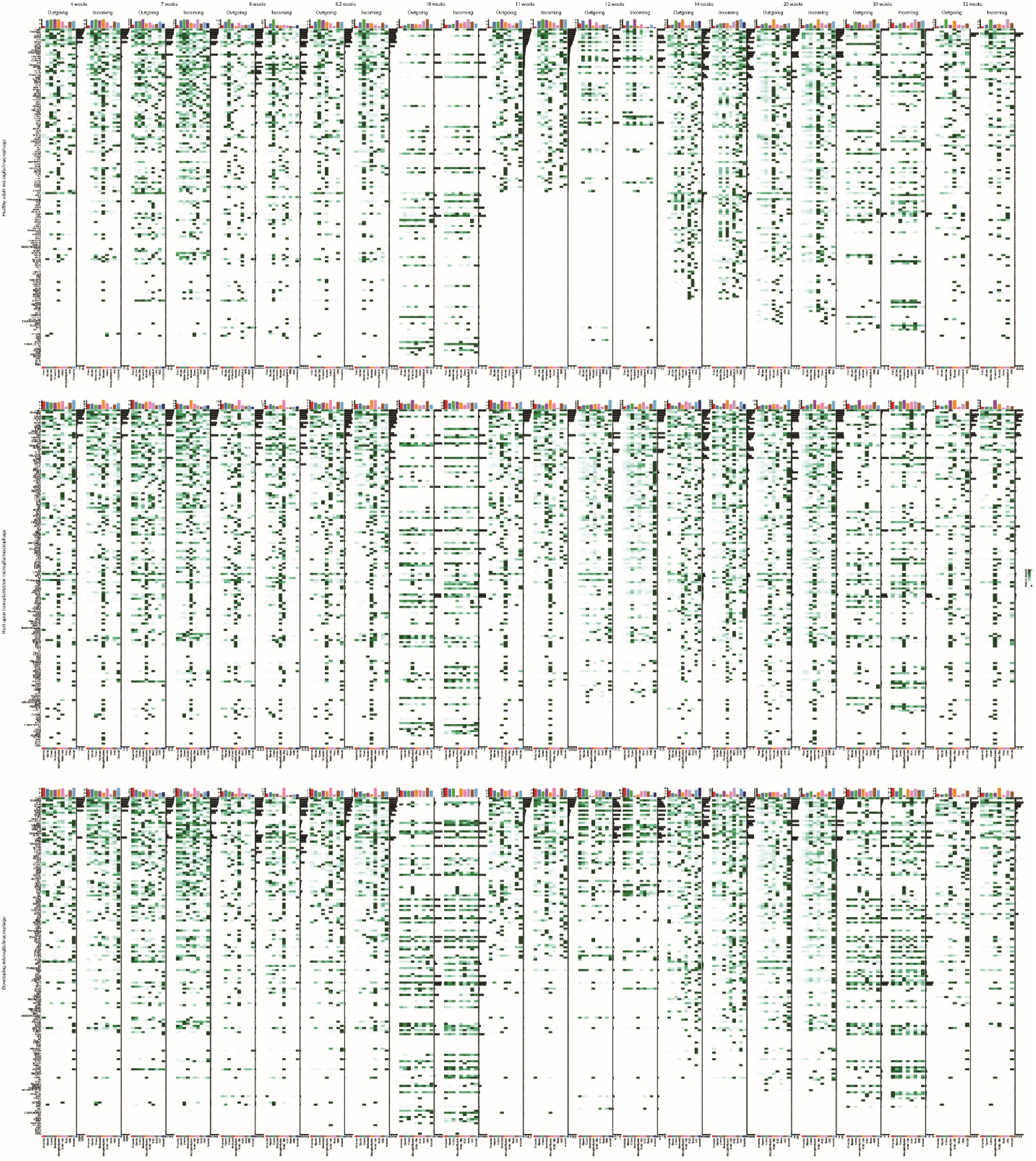
Cell-cell interactions panels for the in silico experiment. Three sets of panels represent three different microglia/macrophage subsets used in the mouse healthy adult retina environment: top – native microglia/macrophage from the healthy adult retina atlas, middle – host upon transplantation microglia/macrophage, bottom – developing microglia/macrophage.

**Figure S4.**
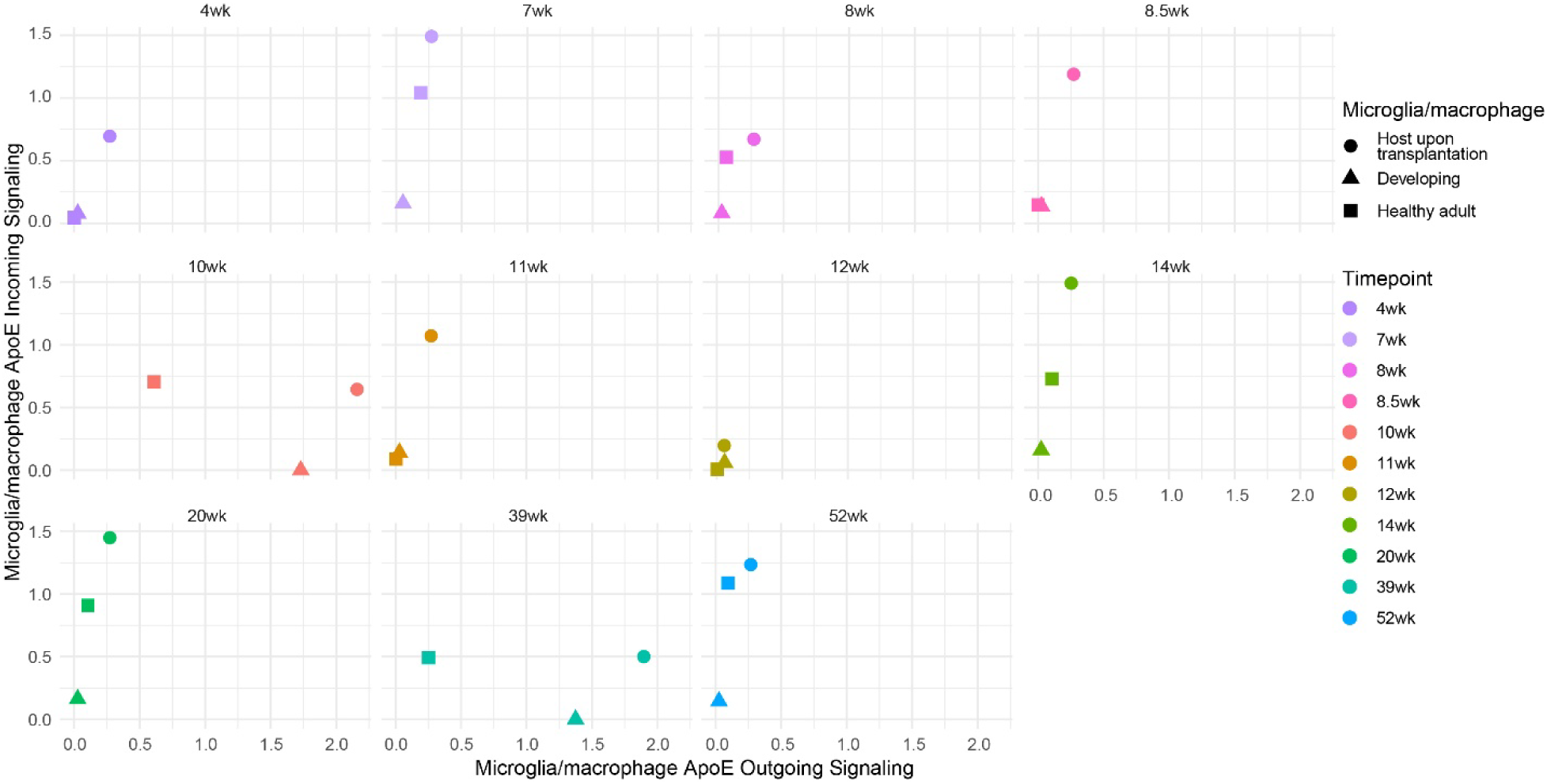
Apoe signaling strength in cell-cell interactions analysis of three sets of microglia/macrophage.

**Figure S5.**
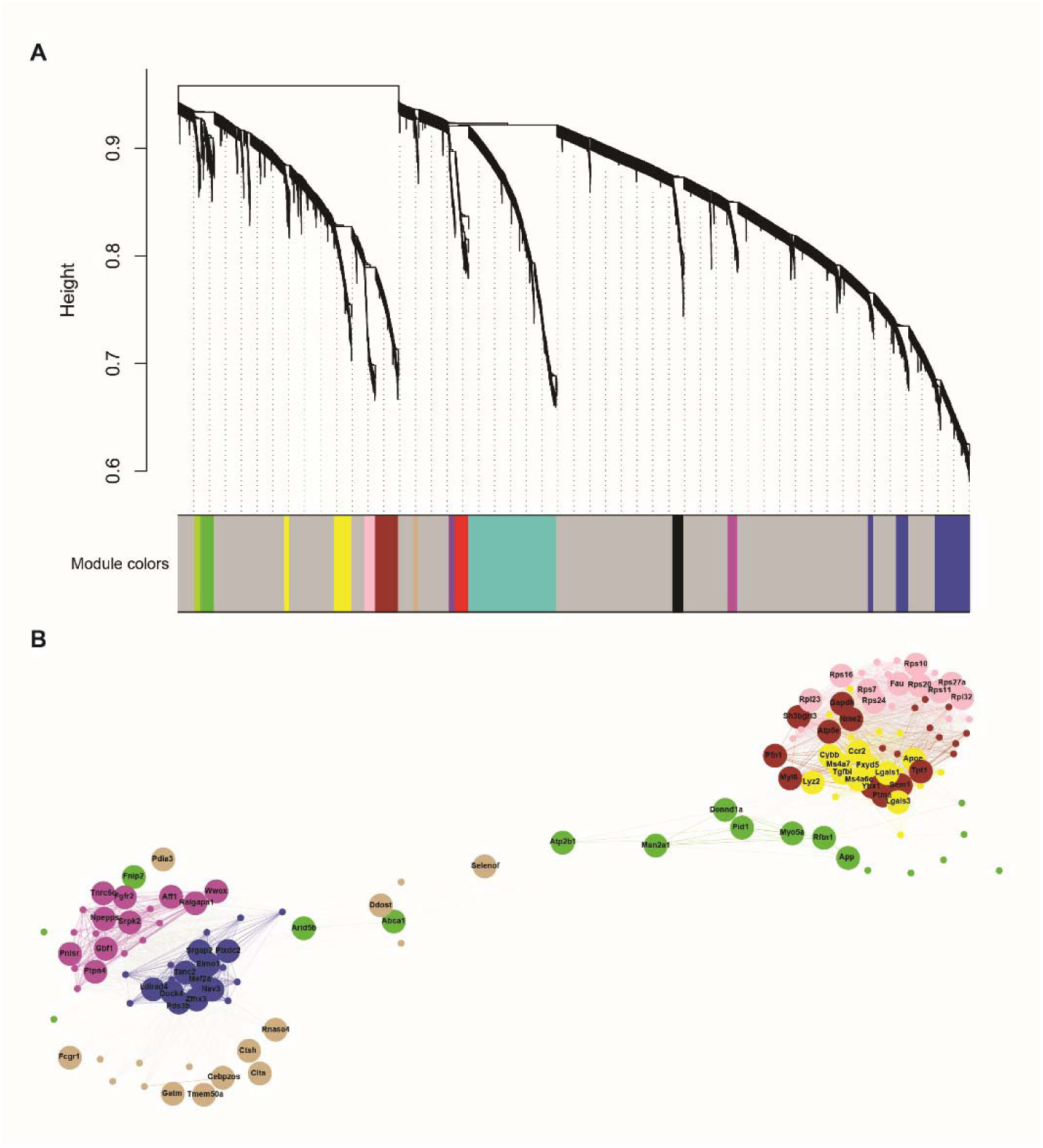
WGCNA analysis of integrated myeloid dataset. **(A)** WGCNA dendrogram demonstrating 12 modules discovered for the integrated myeloid dataset. (B) Network analysis of hub genes in the modules with the highest score in host upon transplantation condition demonstrates the modules clustering into two distinct clusters.

**Figure S6.**
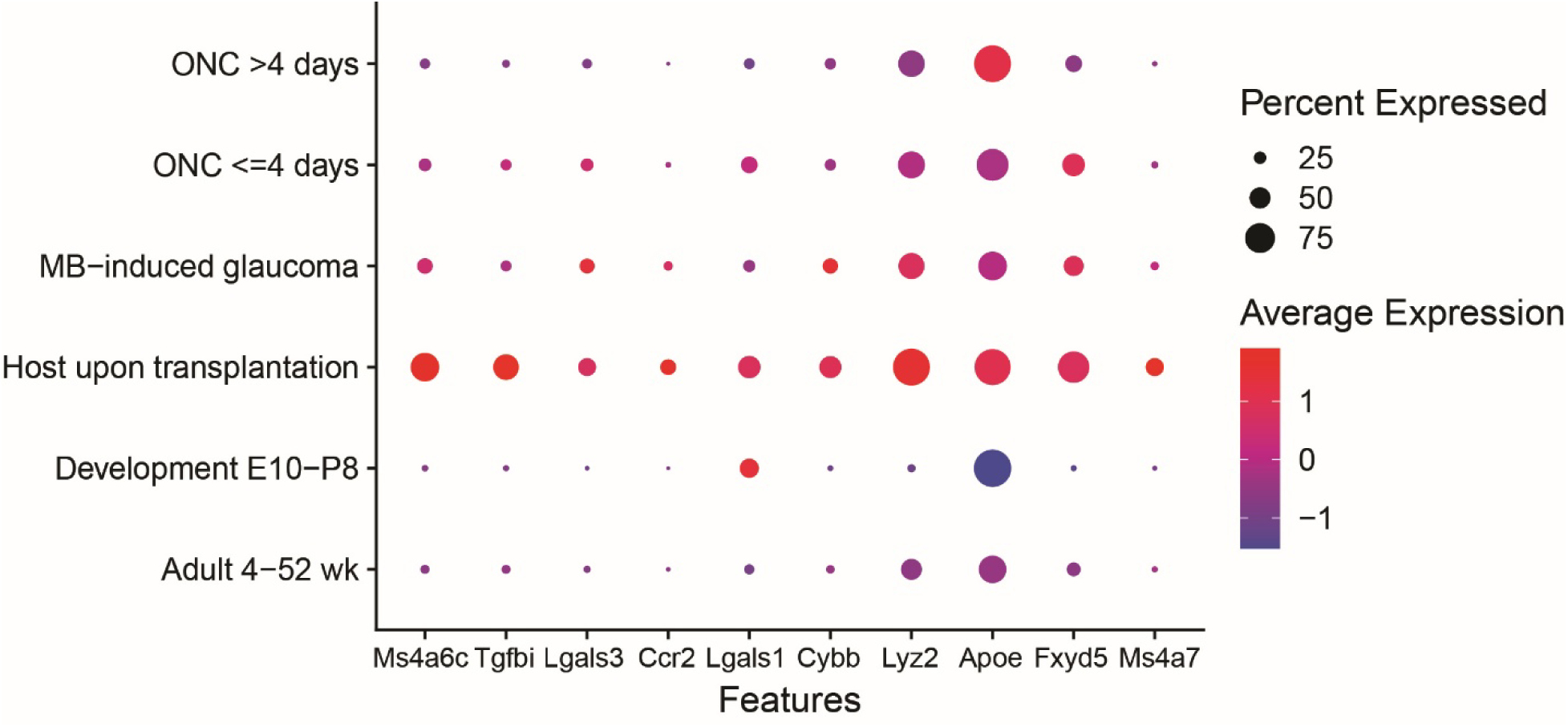
Dotplot demonstrating the expression of top hub genes from HUT11 across the conditions in the integrated myeloid dataset.

**Figure S7.**
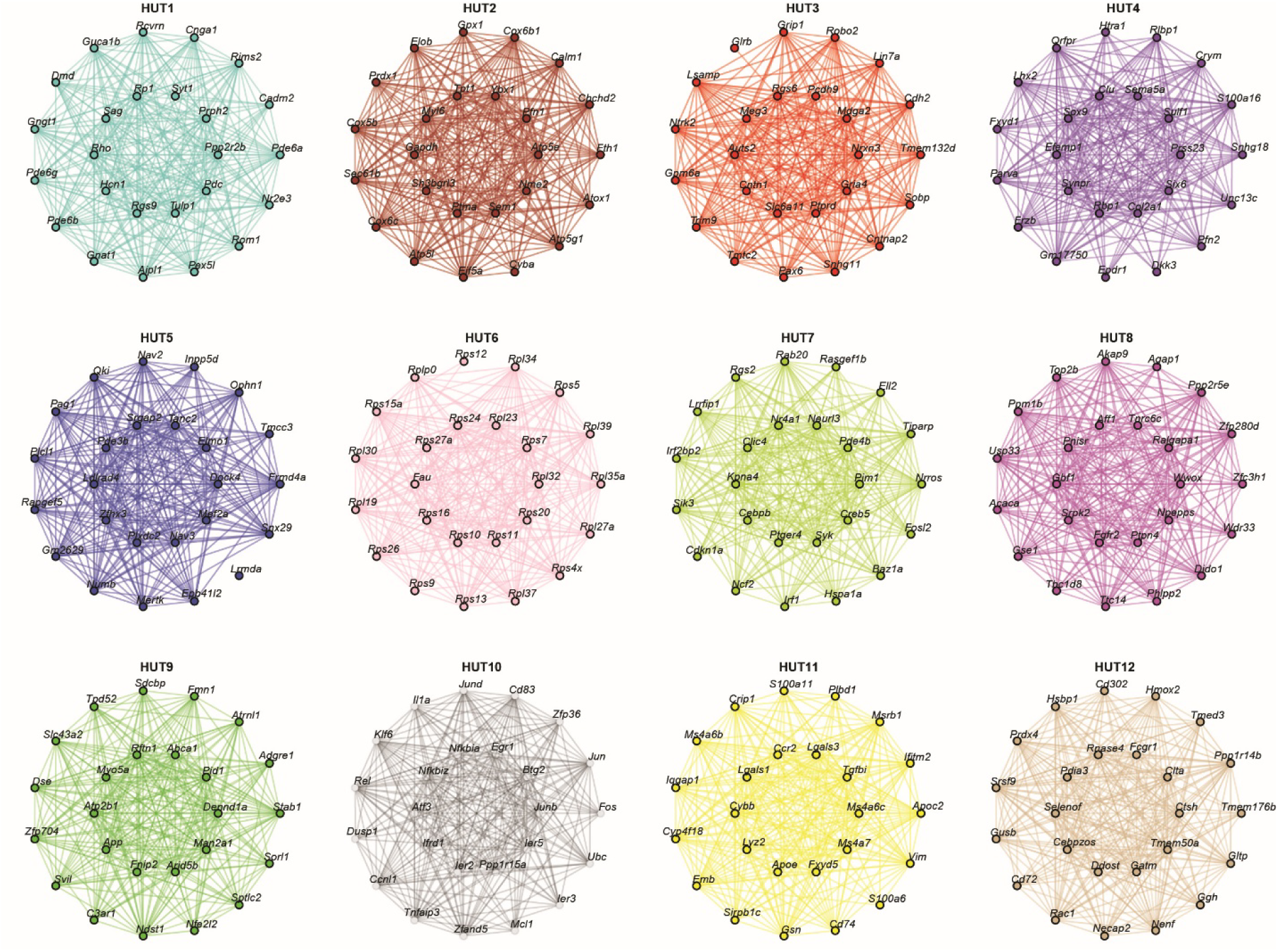
Top 25 genes in each module generated through WGCNA analysis of integrated myeloid dataset.

**Figure S8.**
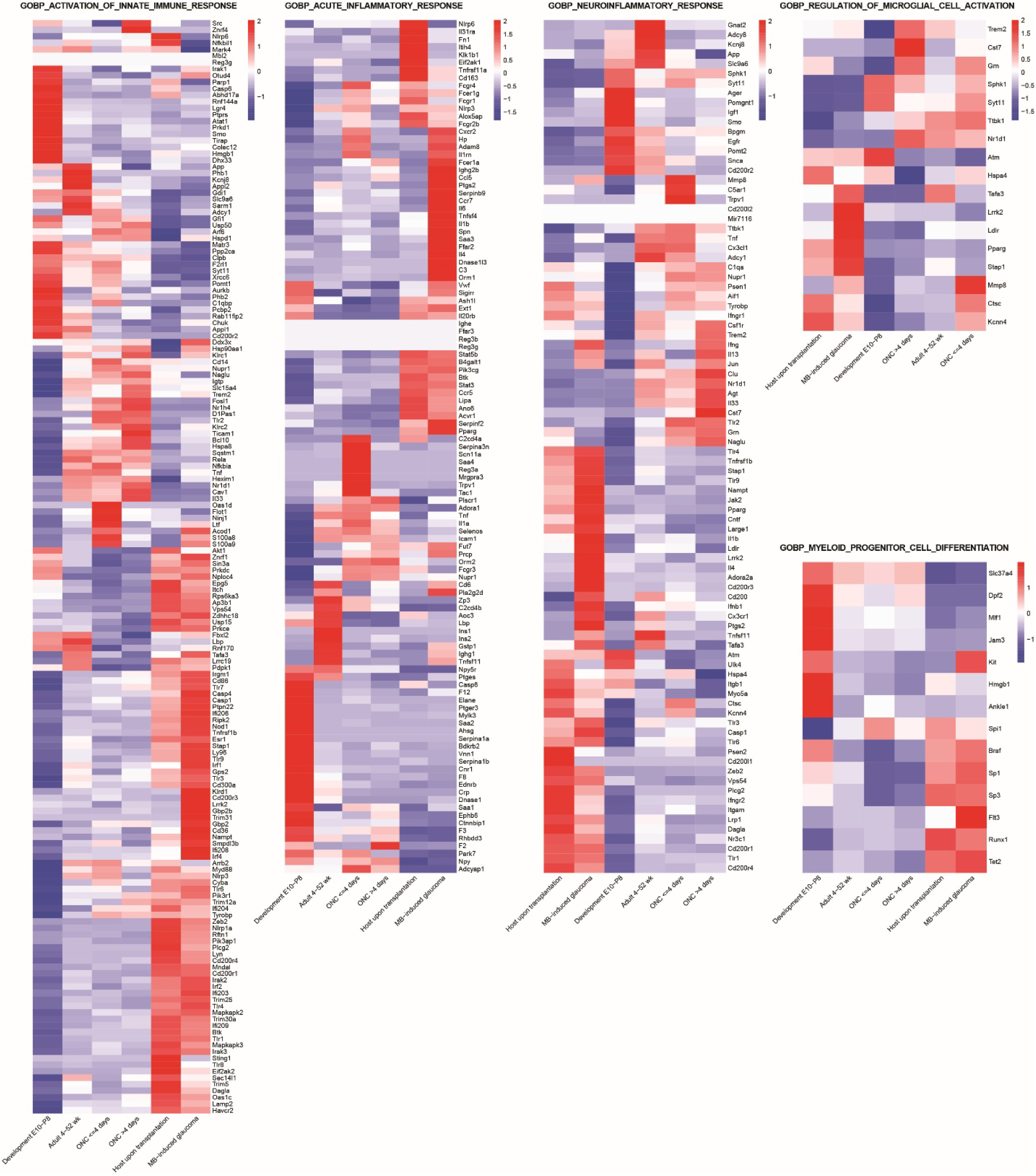
Heatmap highlighting the expression patterns of the genes belonging to the pathways studied in the integrated myeloid dataset.

**Supplementary Table 1.**

| **GSE** | **DOI** | **Condition** |
| --- | --- | --- |
| GSE123758 | Not published_3, as in CELLxGENE | Mouse healthy adult retina atlas |
| GSE125708 | 10.1073/pnas.1821122116<br>(https://pubmed.ncbi.nlm.nih.gov/30988181/) | Mouse healthy adult retina atlas |
| GSE132229 | 10.1073/pnas.1915571116<br>(https://pubmed.ncbi.nlm.nih.gov/31843893/) | Mouse healthy adult retina atlas |
| GSE137398 | 10.1016/j.neuron.2019.11.006<br>(https://pubmed.ncbi.nlm.nih.gov/31784286/) | Mouse healthy adult retina atlas |
| GSE137828 | 10.1016/j.neuron.2019.11.006<br>(https://pubmed.ncbi.nlm.nih.gov/31784286/) | Mouse healthy adult retina atlas |
| GSE153674 | PMID: 33088174<br>(https://pubmed.ncbi.nlm.nih.gov/33088174/) | Mouse healthy adult retina atlas |
| GSE164044 | 10.1016/i.neuron.2019.08.002<br>(https://pubmed.ncbi.nlm.nih.gov/31493975/) | Mouse healthy adult retina atlas |
| GSE169262 | 10.1038/s41467-021-27924-y<br>(https://pubmed.ncbi.nlm.nih.gov/35017532/) | Mouse healthy adult retina atlas |
| GSE176204 | 10.1016/i.celrep.2023.112889<br>(https://pubmed.ncbi.nlm.nih.gov/37527036/) | Mouse healthy adult retina atlas |
| GSE183572 | 10.1038/s42003-022-03676-3<br>(https://www.nature.com/articles/s42003-022-03676-3) | Mouse healthy adult retina atlas |
| GSE199317 | 10.1038/s41590-023-01437-w<br>(https://pubmed.ncbi.nlm.nih.gov/36807640/) | Mouse healthy adult retina atlas |
| GSE204880 | 10.1038/s42003-023-05300-4<br>(https://pubmed.ncbi.nlm.nih.gov/37670124/) | Mouse healthy adult retina atlas |
| GSE205123 | 10.1016/j.ygeno.2023.110644<br>(https://pubmed.ncbi.nlm.nih.gov/37279838/) | Mouse healthy adult retina atlas |
| GSE220626 | 10.3389/fcell.2023.1252547<br>(https://pubmed.ncbi.nlm.nih.gov/37691820/) | Mouse healthy adult retina atlas |
| GSE232470 | 10.1186/s40478-023-01580-3<br>(https://pubmed.ncbi.nlm.nih.gov/37226256/) | Mouse healthy adult retina atlas |
| GSE236389 | Not published_1, as in CELLxGENE | Mouse healthy adult retina atlas |
| GSE239347 | 10.1073/pnas.2314698120<br>(https://pubmed.ncbi.nlm.nih.gov/38064509/) | Mouse healthy adult retina atlas |
| GSE243100 | 10.2337/db23-0368<br>(https://pubmed.ncbi.nlm.nih.gov/37976214/) | Mouse healthy adult retina atlas |
| GSE243413 | 10.1016/i.isci.2024.109916<br>(https://pubmed.ncbi.nlm.nih.gov/38812536/) | Mouse healthy adult retina atlas |
| GSE251717 | 10.1101/2024.03.18.585452<br>(https://pubmed.ncbi.nlm.nih.gov/38562864/) | Mouse healthy adult retina atlas |
| GSE263877 | 10.1038/s41467-024-50033-5<br>(https://pubmed.ncbi.nlm.nih.gov/39009597/) | Mouse healthy adult retina atlas |
| GSE137398 | 10.1016/i.neuron.2019.11.006<br>(https://pubmed.ncbi.nlm.nih.gov/31784286/) | Mouse optic nerve crush retina |
| GSE199317 | 10.1038/s41590-023-01437-w<br>(https://pubmed.ncbi.nlm.nih.gov/36807640/) | Mouse optic nerve crush retina |
| GSE201254 | 10.1016/i.neuron.2022.06.002<br>(https://pubmed.ncbi.nlm.nih.gov/35767994/) | Mouse optic nerve<br>crush retina |
| GSE206622 | 10.1016/j.neuron.2022.06.022<br>(https://pubmed.ncbi.nlm.nih.gov/35952672/) | Mouse optic nerve<br>crush retina |
| GSE206623 | 10.1016/j.neuron.2022.06.022<br>(https://pubmed.ncbi.nlm.nih.gov/35952672/) | Mouse optic nerve<br>crush retina |
| GSE206624 | 10.1016/j.neuron.2022.06.022<br>(https://pubmed.ncbi.nlm.nih.gov/35952672/) | Mouse optic nerve<br>crush retina |
| GSE206625 | 10.1016/j.neuron.2022.06.022<br>(https://pubmed.ncbi.nlm.nih.gov/35952672/) | Mouse optic nerve<br>crush retina |
| GSE223530 | 10.1073/pnas.2305947121<br>(https://pubmed.ncbi.nlm.nih.gov/38289952/) | Mouse optic nerve<br>crush retina |
| GSE228627 | 10.1101/2023.11.17.567614<br>(https://pubmed.ncbi.nlm.nih.gov/38014168/) | Mouse optic nerve<br>crush retina |
| GSE232470 | 10.1186/s40478-023-01580-3<br>(https://pubmed.ncbi.nlm.nih.gov/37226256/) | Mouse optic nerve<br>crush retina |
| GSE241268 | 10.1038/s41467-024-46428-z<br>(https://pubmed.ncbi.nlm.nih.gov/38467611/) | Mouse optic nerve<br>crush retina |
| GSE248537 | 10.1038/s41467-024-46428-z<br>(https://pubmed.ncbi.nlm.nih.gov/38467611/) | Mouse optic nerve<br>crush retina |
| GSE226456 | 10.1016/i.celrep.2023.112889<br>(https://pubmed.ncbi.nlm.nih.gov/37527036/) | Mouse MB-induced<br>glaucoma retina |
| GSE247776 | 10.1038/s41467-024-50050-4<br>(https://pubmed.ncbi.nlm.nih.gov/39080269/) | Mouse MB-induced<br>glaucoma retina |
| not published | not published, as in CELLxGENE (Dong<br>Feng Chen Lab) | Mouse MB-induced<br>glaucoma retina |
| GSE118614 | 10.1016/i.neuron.2019.04.010<br>(https://pubmed.ncbi.nlm.nih.gov/31128945/) | Mouse healthy<br>developing retina |
| GSE122466 | 10.1242/dev.178103<br>(https://pubmed.ncbi.nlm.nih.gov/31399471/) | Mouse healthy<br>developing retina |
| GSE139904 | 10.1126/sciadv.abj9846<br>(https://pubmed.ncbi.nlm.nih.gov/34757798/) | Mouse healthy<br>developing retina |
| GSE186069 | 10.1038/s41586-024-07069-w<br>(https://pubmed.ncbi.nlm.nih.gov/38947520/) | Mouse healthy<br>developing retina |
| GSE172101 | 10.1242/dev.199399<br>(https://pubmed.ncbi.nlm.nih.gov/34143204/) | Mouse healthy<br>developing retina |
| GSE202734 | 10.1016/j.celrep.2023.112237<br>(https://pubmed.ncbi.nlm.nih.gov/36924502/) | Mouse healthy<br>developing retina |
| GSE222956 | 10.1016/j.celrep.2023.112237<br>(https://pubmed.ncbi.nlm.nih.gov/36924502/) | Mouse healthy<br>developing retina |
| GSE228590 | 10.1038/s41586-024-07069-w<br>(https://pubmed.ncbi.nlm.nih.gov/38947520/) | Mouse healthy<br>developing retina |
| GSE245542 | 10.1016/i.celrep.2024.114291<br>(https://pubmed.ncbi.nlm.nih.gov/38823017/) | Mouse healthy<br>developing retina |
| GSE252861 | 10.1016/i.isci.2024.110100<br>(https://pubmed.ncbi.nlm.nih.gov/38947520/) | Mouse healthy<br>developing retina |

